# Conjunction of Factors Triggering Waves of Seasonal Influenza

**DOI:** 10.1101/168476

**Authors:** Ishanu Chattopadhyay, Emre Kıcıman, Joshua W. Elliott, Jeffrey L. Shaman, Andrey Rzhetsky

## Abstract

Understanding the subtle confluence of factors triggering pan-continental, seasonal epidemics of influenza-like illness is an extremely important problem, with the potential to save tens of thousands of lives and billions of dollars every year in the US alone. Beginning with several large, longitudinal datasets on putative factors and clinical data on the disease and health status of over 150 million human subjects observed over a decade, we investigated the source and the mechanistic triggers of epidemics. Our analysis included insurance claims for a significant cross-section of the US population in the past decade, human movement patterns inferred from billions of tweets, whole-US weekly weather data covering the same time span as the medical records, data on vaccination coverage over the same period, and sequence variations of key viral proteins. We also explicitly accounted for the spatio-temporal auto-correlations of infectious waves, and a host of socioeconomic and demographic factors. We carried out multiple orthogonal statistical analyses on these diverse, large geo-temporal datasets to bolster and corroborate our findings. We conclude that the initiation of a pan-continental influenza wave emerges from the simultaneous realization of a complex set of conditions, the strongest predictor groups are as follows, ranked by importance: (1) the host population’s socio- and ethno-demographic properties; (2) weather variables pertaining to relevant area specific humidity, temperature, and solar radiation; (3) the virus’ antigenic drift over time; (4) the host population’s land-based travel habits, and; (5) the spatio-temporal dynamics’ immediate history, as reflected in the influenza wave autocorrelation. The models we infer are demonstrably predictive (area under the Receiver Operating Characteristic curve ≈ 80%) when tested with out-of-sample data, opening the door to the potential formulation of new population-level intervention and mitigation policies.

## Introduction

Seasonal influenza is serious threat to public health, claiming tens of thousands of lives every year. A large number of past studies focus on identifying the likely factors responsible for initiating each seasonal disease wave. Typically, each such study focuses on one or a few hypothetical factors. Our study aimed at an integrative joint analysis of numerous suggested disease triggers, comparing their relative importance and possible cooperation in triggering pan-US waves of seasonal influenza infection.

The traditional empirical approach of testing a causal link between a factor and an outcome of an experiment was to vary one factor at a time, while keeping the other factors (experimental conditions) constant. This “all the rest of conditions is equal” assumption is often referred to by its Latin form as *ceteris paribus.* R.A. Fisher ([19], p. 18) noted that, in real-life experiments, perfect *ceteris paribus* is not achievable “because uncontrollable causes which may influence the results are always … innumerable.” Fisher’s proposed solution to this problem is to design experiments to involve random assignment of treatment (the putative causal factors states) to individual experiments and then use regression analysis to estimate the value and significance of the putative causal effect.

Because designing a true randomized experiment studying infection propagation in humans is not ethical, the next best alternative is to perform observational studies in which human infection data is generated in the absence of the investigators control, by passive collection of large volumes of health statistics. Under certain conditions, an observational study can provide an approximation to randomized experiment [[21], p. 226]: “In a randomized experiment, the groups receiving each treatment will be similar, on average, or with differences that are known ahead of time by the experimenter (if a design is used that involves unequal probabilities of assignment). In an observational study, in contrast, the groups receiving each treatment can differ greatly, and by factors not measured in the study. A well-conducted observational study can provide a good basis for drawing causal inferences, provided that it: (1) controls well for background differences between the units exposed to the different treatments; (2) has enough independent units exposed to each treatment to provide reliable results (that is, narrow-enough posterior intervals); (3) is designed without reference to the outcome of the analysis; (4) minimizes or appropriately models attrition, dropout, and other forms of unintentional missing data, and; (5) takes into account the information used in its design.”

Hypothesis-driven science, wherein investigators formulate a single, testable hypothesis and design specific experiments to test it, is a core element of the scientific method, and works well in most scientific fields. However, a new challenge emerges in data-rich scientific fields, such as genomics, epidemiology, economics, climate modeling, and astronomy: How do we choose the most promising hypotheses among millions of eligible candidates that potentially fit data? One solution to this challenge is the many-hypotheses approach, a method of automated hypothesis generation in which many hypotheses are systematically produced and simultaneously tested against all available data. This approach is currently used, for example, in whole-genome association or genetic linkage studies, and often enables truly unexpected discoveries. In contrast to the single-hypothesis approach, the many-hypotheses approach explicitly accounts for the large universe of possible hypotheses through calibrated statistical tests, effectively reducing the likelihood of accidentally accepting a mediocre hypothesis as a proxy for the truth [45].

The many-hypotheses computation approach provides a complement to carefully controlled and highly focused wet laboratory experiments. Running controlled experiments to test a single hypothesis necessarily ignores many of the complexities of a real-world phenomenon; these complexities are necessarily present in large, longitudinal datasets. Of course, the data-driven “many-hypotheses” approach is only one aspect of the broader scientific process progressing toward the development of verifiable general theories.

As all causality detection methods come with dissimilar limitations and are imperfect in unique ways, we designed our study intentionally to attack the same target problem using three different statistical approaches, shown in Figure 1 A: **Approach 1:** A non-parametric Granger analysis [27] focusing on infection flows’ directionality across the US and whether influenza propagates via long- vs. short-distance travel (we run analysis across all pairs of air- and land-travel county neighbors, respectively); **Approach 2:** A mixed-effect Poisson regression [31] explicitly accounting for auto-correlation of the infection waves in time and space, along with the full set of socioeconomic, climate, and geographic predictors, and; **Approach 3:** A county-matching, nonparametric analysis to identify the minimum predictive set of factors that distinguish counties associated the onset of influenza season season [43].

**Fig. 1.**
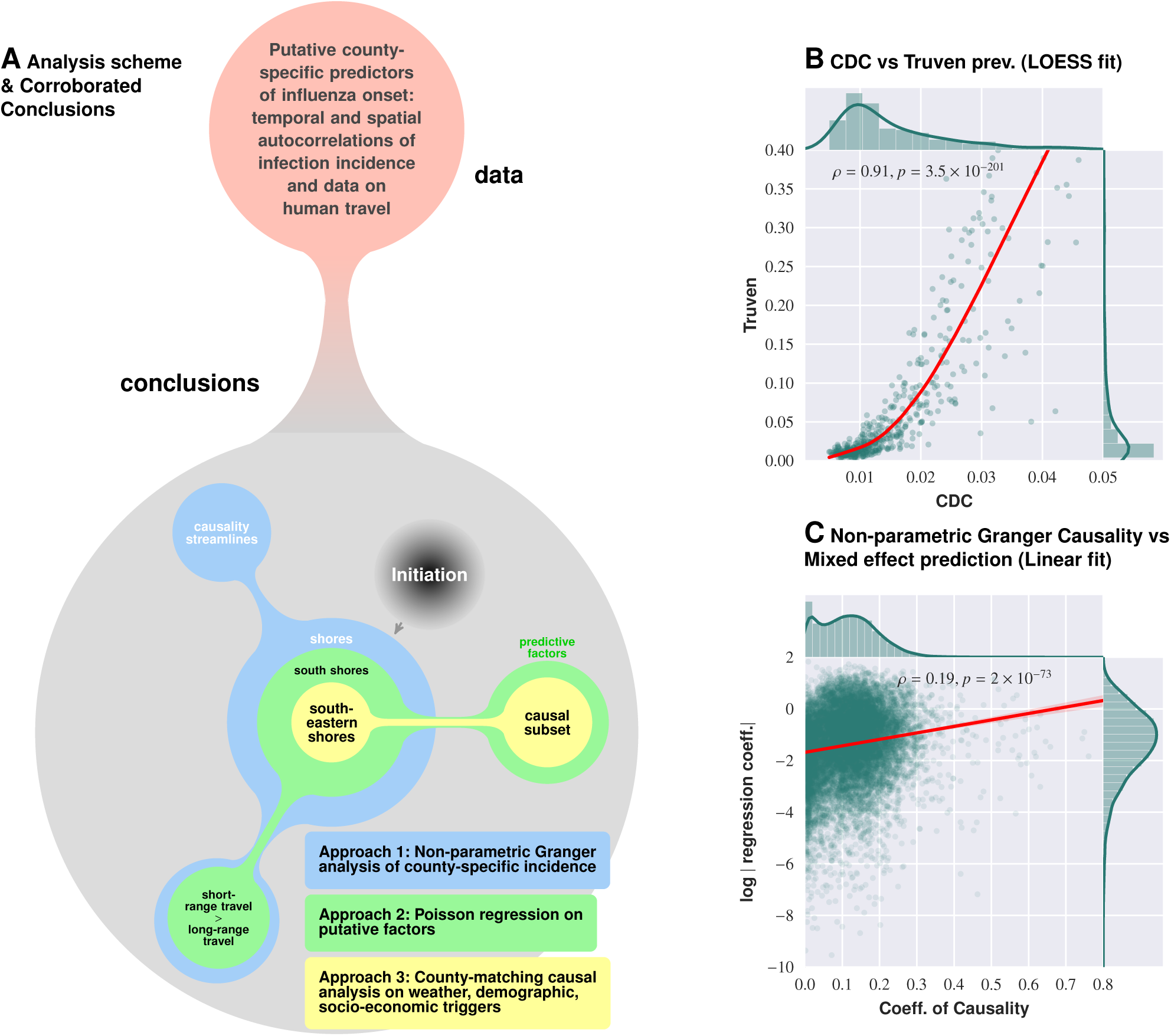
Logical flow and cross-corroboration of conclusions. Plate A: Diverse data sets processed via multiple techniques to reach convergent, and reinforcing, conclusions. Approach 1 (Granger-causality analysis) shows that the epidemic tends to begin near water-bodies, and that short-range travel is more influential compared to air-travel for propagation. Approach 2 (Poisson regression) identifies significant predictive factors, suggesting that the epidemic begins near the southern shores of the US, and corroborates the result on short- vs. long-range travel. Approach 3 (county-matching) points to south eastern shores of the continental US as where the epidemic initiates, and identifies a validated subset of predictive factors. Plate B shows influenza prevalence as reported by Truven data set positively correlates with CDC reports. Plate C illustrates that our conclusions regarding a putatively causal influence between neighboring counties, inferred using different techniques (mixed-effect regression vs. non-parametric Granger-causality), match up positively.

Our study became possible through access to several, very large longitudinal datasets: (1) a nine-year collection of insurance records capturing the dynamics of influenza-like illnesses (ILIs) in the United States; (2) temporally collinear, high-resolution weather measurements over every US county; (3) detailed air travel [56] and geographic proximity data [57] showing connectivity between US counties; (4) billions of geo-located Twitter messages reflecting long- and short-distance human movement patterns, and; (5) US census data accounting for US county and county-equivalent population distribution, demographic, and socioeconomic properties [33]. We also used influenza-like illnesses (ILIs) from the whole-country insurance claims (Truven MarketScan database, see Methods). An explicit comparison of ILIs in the insurance claims to influenza records provided by the Center for Disease Control and Prevention [9] showed that the two sources agree well (ρ = 0.91, p = 3.5 × 10^−201^), with insurance claims providing higher data resolution, see Figure 1 B. Curiously, the relationship between the two sources of ILIs observations is not linearly related: We attribute this to the lower resolution of the CDC data. These three types of analysis produce congruent–albeit not identical–results. (For example, Figure 1 C shows comparisons of putative influence parameters produced by Granger-causality analysis with conventional regression coefficients.)

## Methods

### Candidate factors in influenza initiation

To investigate county-specific variability, we group candidate factors into several categories: demographic, relation to human movement, and climatic.

#### Demographic

Influenza is transmitted through direct contact with infected individuals, via contaminated objects, and via virus-laden aerosols. Thus, human population density (how many people happen to be around?) and social connectivity (how many people interact with each other and how frequently [5]?) are factors expected to affect local virus incubation and spread. In addition to population density, we consider socioeconomic factors such as mean household income, levels of poverty and urbanity, as well as the prevalence of ethnic and age groups (see Figure 2).

**Fig. 2.**
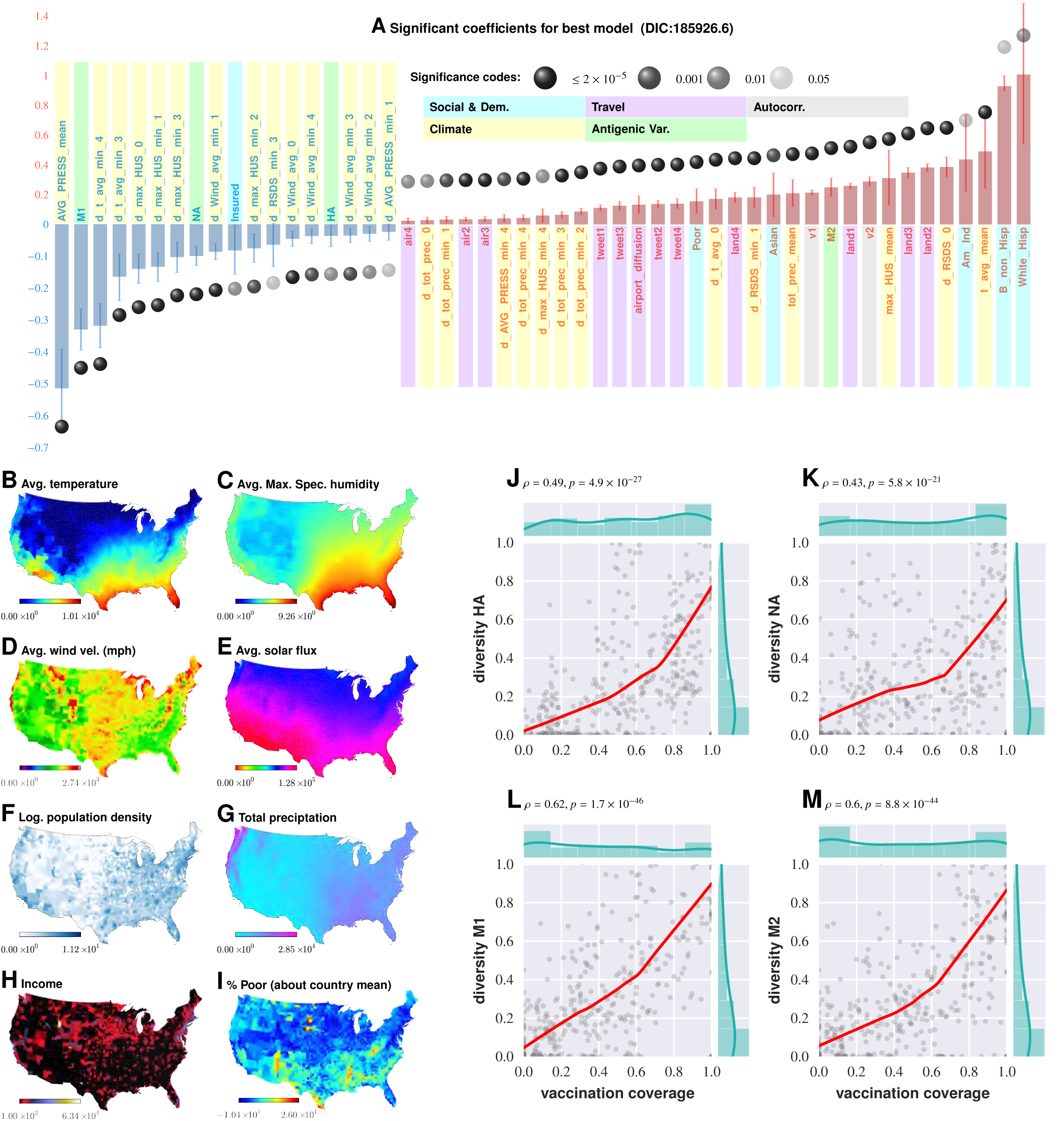
Putative determinants of seasonal influenza in the continental US and Poisson mixed-effect regression analysis (Approach 2). Plate A shows the significant variables along with their computed influence coefficients from the mixed-effect Poisson regression analysis (best model chosen from 126 different regression equations with different variable combinations). The statistically significant estimates of fixed effects are grouped into several classes: climate variables, economic and demographic variables, autoregression variables, variables related to travel, and those related to antigenic diversity (see the last entry in Table V for the detailed regression equation used. The complete list of all models considered is given in Table ??). Plates B - I enumerate the average spatial distribution of a few key significant factors considered in Poisson regression: (B) Average temperature; (C) Average maximum specific humidity; (D) Average wind velocity in miles per hour; (E) Average solar flux; (F) Logarithm of population density (people per square mile); (G) Total precipitation; (H) Income, and (I) Percent of poor as deviations about the country average. Plates J-M show the strong dependence between our estimated antigenic diversity corresponding to the HA, NA, M1, and M2 viral proteins, and cumulative fraction of inoculated population, where both sets of variables are geo-spatially and temporally stratified.

#### Human movement

We consider two measurements of human movement: 1) The first measurement reflects the proximity of counties to major airports. We compute an exponentially-diffusing influence from counties with major US airports, weighted by passenger enplanements at their respective locations. This accounts for people moving to and from both major airports and neighboring counties. 2) The second measurement is a large-scale movement matrix representing people’s week-to-week travels between counties. These data are culled from a complete collection of geo-located Twitter messages, captured over 3.5 years, and constitutes a large-scale, longitudinal sample of individual movements. We used only automatically geo-tagged tweets.

We observe patterns consistent with both intuitive patterns and prior studies of large-scale movements; the majority of people in a county remain in the same county from week-to-week, with people traveling to metropolitan areas in proportion to their size and in inverse proportion to their distance. These data compare favorably with other publicly available datasets on human mobility. Admittedly, there is a bias in the Twitter movement data, with younger, urban, and higher-income populations more likely than older, rural, and lower-income populations to use social media technology [49]. However, bias is a problem associated with any data on human mobility.

#### Climate variables

Specific humidity and a drop in temperature have been suggested as the key drivers in triggering seasonal influenza epidemics [40], [51], [52]. These initial conclusions were drawn from experiments conducted using an animal model (guinea pig), under controlled laboratory conditions [51], [40], followed by indirect support from epidemiological modeling [52].

#### Antigenic Variation

The influenza virus counteracts host immunity via subtle genomic changes over time. The more gradual process, known as antigenic drift, is a manifestation of a gradual accumulation of mutations within viral surface proteins recognized by human antibodies, such as hemagglutinin (HA) and neuraminidase (NA). These mutations are typically introduced during cycles of viral replication [7]. Most of these mutations are neutral, i.e. they do not affect the functional conformation of the viral proteins. However, some of these alterations do in fact change secondary and tertiary protein structures sufficiently to have a negative impact on the binding of host antibodies raised in response to previously circulating strains [61]. (Many of such mutations also reduce viability of the virus.) Thus, while a single infection episode is potentially enough to provide long-term host immunity to the invading strain, antigenic variation due to intense selection pressure gives rise to novel viral strains, making re-infections possible within the span of a few years [2]. This kind of perpetual Red-Queen arms race injects into influenza dynamics auto-correlative dependencies over multiple seasons. It has been suggested that substantial antigenic drift might be associated with more severe, early-onset influenza epidemic, resulting in increased mortality [58]. In contrast to antigenic drift, *antigenic shift* is an abrupt, major change in virus structure due to gene segment re-assortment that occurs during simultaneous infection of a single host by multiple influenza subtypes [15]. Antigenic shift results in new versions of viral surface proteins. Antigenic shift due to re-assortment give rise to novel influenza subtypes that, if capable of sustained human-to-human transmission, can have devastating consequences for the human populations, e.g. the 2009 H1N1 pandemic [44].

We factor in the potential effect of antigenic variation in our analysis by estimating the surface protein’s population diversity – hemagglutinin (HA), neuraminidase (NA), matrix protein 1 (M1), and matrix protien M2 – as a function of time and geographical sample collection location. Our rationale for our focus on these proteins is that HA, NA, and to some degree M1, are present on the viral surface [37], contribute to viral assembly, and mediate the release of membrane-enveloped particles [11]. M2 has been shown to have enhanced the pandemic 2009 influenza A virus [(H1N1)pdm09] [20] HA-pseudovirus infectivity [1]. We find that antigenic diversity in all four of the viral proteins we considered are significant predictors.

Interestingly, while the increasing diversity found in HA, NA, and M1 inhibits the epidemic trigger, the higher diversity in M2 enhances it (see Discussion).

#### Vaccination Coverage

Vaccination is widely regarded as our most promising tool to combat influenza, though antigenic variation between seasons makes it difficult to craft an effective vaccination strategy [6]. Understanding how the virus will evolve in the short-term is key to finding the correct antigenic match for an upcoming influenza season. Additionally, short-term molecular evolution might rapidly give rise to immune-escape variants that, if detected, might dictate intra-season updates in vaccine composition. More importantly, vaccination itself might exert significant selection pressure to influence antigenic drift. The effect of vaccination on viral evolution has been documented in an avian H5N2 lineage in vaccinated chickens [39], suggesting that similar processes might be occurring in human counterparts. The diversity of the surface proteins at any point in time between seasons suggests that our current vaccination strategies are limited to confer partial immunity, which can result in a highly immune or vaccinated population selectively pressurizing the viral population to evolve more quickly than usual. Given that influenza moves quickly across geographies and there are multiple co-circulating strains that may confer partial cross-protection, it is a moving target. Further, there are complexities due to early-life imprinting (original antigenic sin). The second clause may not be common given this imprinting and the variability of the efficacy of the vaccine (often across demographic strata).

We factor in the effect of vaccination coverage by estimating the cumulative fraction of the population that received the current influenza vaccine stratified by geo-spatial location and time of inoculation within each influenza season. Our analysis indicates that vaccination coverage is not a significant predictor–at least in our best Poisson regression model (model quality measured by Deviance Information Criterion), which might be a consequence of the observed dependence between vaccination coverage and antigenic diversity (see Figure 2, Plates J-M).

#### Return to School Effect

Social contact among children in schools has been extensively investigated as a determinant of the peak incidence rate. This is one of the few factors that might lend itself to intervention relatively easily, and hence the interest is well-justified. While any reduction in social contact should, in theory, directly impact transmission, quantifying the effect of this specific mode of contact on the incidence rate has been difficult to calculate. Predictions of the reduction in the peak incidence associated with reduced social contact were typically 20 - 60% [18], [29], with some studies predicting much larger reductions of ≧ 90% [42], [22]. Reductions in the cumulative attack rate (AR, ratio of the number of new cases to the size of the population at risk) were usually smaller than those in the peak incidence. Several studies predicted small (~10%) or no reduction in the cumulative AR [18], [29], [64], [16], [63], [36], [14], [50], [60], [23], [38], [62], [65], whilst a few predicted substantial reductions (e.g. ≧ 90%) [24], [14], [17], [13]. Only two studies [24], [38] predicted that peak incidence might increase markedly under some circumstances following school closures, e.g. by 27% if school closures caused a doubling in the number of contacts in the household and community, or by 13% if school systems were closed for two weeks at a prevalence of 1% in the general population. Studies have also investigated the effect of such interventions on children vs. adults; one study predicted an overall reduction in the cumulative AR, but an increase of up to 48% in the cumulative AR for adults in some situations [3]. While this diverse set of predictions in the literature often pertains to the effect of school closures as an intervention tool, we are more interested in the influence that the current school schedule has, if any, on triggering the epidemic. To answer this specific question, we formulated a simple statistical test to determine whether the timing of return-to-school after summer and winter holidays significantly predicts influenza season initiation. We found insufficient evidence in support of this effect (see Methods & Materials).

### Clinical data source

The source of the clinical incidence data used in this study is the Truven Health Analytics MarketScan^®^ Commercial Claims and Encounters Database for the years 2003 to 2012. The database consists of approved commercial health insurance claims for between 17.5 and 45.2 million people annually, with linkage across years, for a total of approximately 150 million individuals. This US national database contains data contributed by over 150 insurance carriers and large, self-insuring companies. We scanned 4.6 billion inpatient and outpatient service claims and identified almost six billion diagnosis codes. After un-duplication, we identified approximately 12.8 unique diagnostic codes per individual. We processed the Truven database to obtain the reported weekly number of influenza cases over a period of 471 weeks spanning from January 2003 to December 2013, at the spatial resolution of US counties. To define influenza in insurance claims, we used the following set of ICD9 codes: 487.8, 488.12, 488.1, 488.0, 488.01, 488.02, 487.0, 487.1, 488.19, 488.09, 488, 487, and 488.11.

### Data on antigenic drift for Influenza A

Sequence data for this computation was obtained from the National Institute of Allergy and Infectious Diseases (NIAID) Influenza Research Database (IRD) [66] through their web site at http://www.fludb.org.

### Data on vaccination coverage

Data on vaccinations is extracted from our EHR database corresponding to the procedural codes 90661 and Q2037, which correspond to the dominant influenza vaccines. (http://flu.seqirus.com/files/billing_and_coding_guide.pdf)

### Data on human air travel

We use a complete, directed graph of passenger air travel for 2010, accounting for the number of passengers transported in each direction [56]. For each county, we compute an air neighborhood network: For counties i and j, the incoming edge to county i represents the proportion of passengers, and p_ji_ represents the ratio of all passengers who traveled from county i to county j by plane, to the total number of travelers who left county i by plane during the year, so that ∑_j≠i_ P_ji_ = 1.

### Data on general human travel patterns in the US

Using the complete Twitter dataset, we aggregate a movement matrix to capture people’s week-to-week travels between counties from geolocated Twitter messages (captured during the period of January 1, 2011 through June 30, 2014). This dataset includes approximately 1.7 × 10^9^ messages and represents 3 × 10^8^ user-days of location information. A small, but significant, percentage of Twitter messages are automatically tagged with the author’s current latitude/longitude information, as determined by their mobile device. Each latitude/longitude-annotated tweet was mapped to a FIPS county code based on Tiger/Line shape files from the 2013 Census dataset (http://www.census.gov/geo/maps-data/data/tiger-line.html). In addition, we calculated a variant of our movement matrix to capture seasonality and other temporal dynamics: A set of 52 movement matrices captured weekly, county-to-county movements, observed in each week of the year, aggregating each week based on the corresponding observations, from each year from 2011 through 2014.

**TABLE I.** Social connectivity: The US southern region appears to have an unusually high level of social connectivity. (In GSS survey results, the number of close friends, close friends who are neighbors, and number of friends who all or mostly know each other is higher in the South, especially in the East/South/Central census region, than in the country at large.)

|  | WSC (TX, OK, AR, LA) | ESC (MS, AL, TN, KY) | SA (FL, GA, SC, NC, VA, WV, MD, DC) | Country-at-large | WNC (ND, SD, NE, KS, MO, IA, MN) (not in South/Southeast) this is the second most social region following ESC |
| --- | --- | --- | --- | --- | --- |
| Close friends | 7.22 | **12.76** | **8.20** | 7.57 | **10.56** |
| Close friends who are neighbors | 1.02 | **3.40** | 1.32 | 1.45 | **3.15** |
| % of friends who all or mostly know each other | All:20% Mostly: 43% | **All:18% Mostly: 58%** | All: 11% Mostly: 52% | All: 12 Mostly: 50% | **All: 16% Mostly: 58%** |
| How often visit closest friends<sup>‡</sup> | 107 | **151** | 126 | 122 | **129** |

<sup>‡</sup>Survey options are: lives in household, daily, several times a week, once a week, once a month, several times a year, and less often. These are converted to approximate number of visits per year (see Supplement for more information about the GSS analysis).

This dataset constitutes a large-scale, longitudinal sample of individual movements. We find that the movement patterns are consistent with intuitive patterns and prior studies of large-scale movements; the majority of people in a county remain in the same county week-over-week. Most travel between counties occurs between neighboring counties, and between counties and large metropolitan areas, conditioned on distance and size of the metropolitan area. We represent our movement data as an n × n matrix **M** that captures the likelihood that a person observed in county i during the course of a week will be observed in county j in the following week. We calculate the entries m_i,j_ of matrix M as follows:

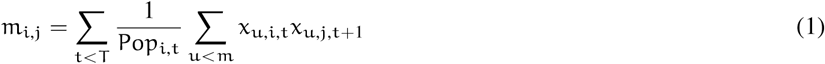

For each time interval, we compute the mean movement vector from a given county by summing all county-to-county movement vectors, weighted by the proportion of people moving in each direction.

To investigate the role of proximity to major airports, we model influence diffusion as follows: Let x_i_ be the i^th^ county, and ν_i_, be the total contribution obtained by diffusing influences from the major airport bearing counties. Let
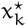
be the k^th^ airport-bearing county, and let N be the total number of major airport locations considered, in our case, N = 27. Let g__k__ be the volume of traffic for the k^th^ major airport location. We then compute v_i_ as follows:

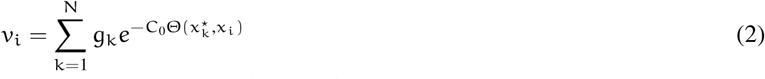

where Θ(·, ·) is the distance in miles between two locations, we computed using the Haversine approximation. The value of the constant C__0__ was chosen to be 0.1. Small variations in the constant do not significantly alter our conclusions. As noted above, proximity to airports had a significant positive influence in sparking seasonal epidemics; the influence is significantly weaker once an outbreak is well under way.

### Estimating antigenic diversity

In this study, we measured antigenic diversity as follows: Let S_i,x,t_ be the set of amino acid sequences for the i^th^ protein (one of HA, NA, M1 or M2), collected in year t (t ranging between 2003 and 2011), in state x of the continental United States. The temporal resolution of the sequence data is thus set to years instead of weeks, and the spatial resolution to states instead of counties. These resolutions are coarse compared to our EHR data on infection incidences, and is set in this manner to maintain sufficient statistical power. For each such set of amino acid sequences S_i,x,t_, we compute the set D(S_i,x,t_) of pairwise edit distances:

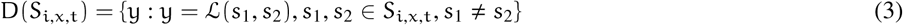

where 𝓛(s_1_, s_2_) is the standard edit distance (also known as the Levenshtein distance) between the sequences s_1_, s_2_. Mathematically, the Levenshtein distance between two strings a, b (of length |a|, |b| respectively) is given by:

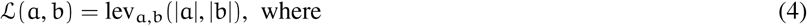

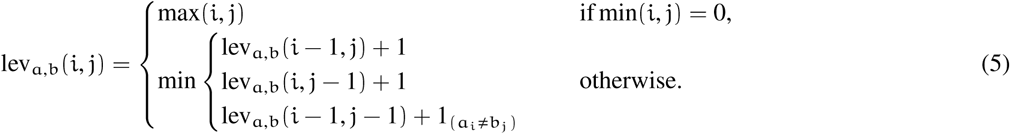

where 1_(a_i_≠b_j_)_ is the indicator function equal to 0 when a_i_ = b_j_ and equal to 1 otherwise, and lev_a,b_(i, j) is the distance between the first i characters of a and the first j characters of b.

The antigenic diversity of the i^th^ protein at time t in state x is then defined as the median of the distribution of the values in the set D(S_i,x,t_). Clearly, as the sequences get more diverse at a point in time and space due to molecular variations brought about by either drifts or shifts, the measure deviates more from zero. Use of the median provides robustness to outliers.

Importantly, here we are not comparing changes in antigenic makeup directly, but estimating the current diversity in the protein primary structure. However, due to our coarse temporal resolution, we expect our measure to be representative of the cumulative changes that occur within each influenza season. State-level geo-spatial variation, and our coverage of 9 epidemic seasons imply that we capture spatio-temporal dependence of population level sequence diversity as a factor influencing the incidence dynamics in individual seasons, which we include as a potential predictor in our Poisson regression analysis.

### Estimating vaccination coverage

We incorporate the effect of vaccination coverage by estimating the cumulative fraction of the population that received the current influenza vaccine within the previous 20 week period. This approach to estimating vaccination coverage does not correct for season-specific antigenic match, or the lack thereof. Nevertheless, because we explicitly include measures of antigenic diversity in addition to vaccination coverage, we expect that effects arising from the degree of antigenic match will indeed be factored in; if there is a significant mismatch, we expect the antigenic diversity in that year to be less and vice verse. This assumes implicitly that vaccination does indeed play a major role in exerting significant selection pressure, an assumption which is justified by our observation of a strong dependency between vaccination coverage and normalized antigenic diversity as shown in Figure 2, Plates J-M.

We find that antigenic diversity is quite strongly affected by vaccination coverage. This reflects the theoretical predictions in Boni et al. [8], where it is shown that the amount of observed antigenic drift increases as immunity in the host population increases and pressures the virus population to evolve.

### Estimating the effect of return-to-school days

The absence of consensus, and the diversity of modeling assumptions pertaining to this effect (described before) makes it difficult to validate the conclusions in large scale epidemiological data. We carried out a simple test to determine if there is statistical evidence that after-holiday return-to-school periods in August-September and in January predict or trigger the seasonal epidemic.

For this we assumed a broad window to cover all such school opening across the continental US including the last week in August, the entire month of September, two weeks in October, and two weeks in January. Then we carried out a Fisher’s exact test to determine if the overlap between these weeks and the identified “trigger period” (See Table ??) for the seasonal epidemic are sufficiently non-random. We ended up with a p-value of 0.84 and an odds ratio of 0.8403, strongly suggesting that school opening dates are not a significant factor in triggering the seasonal epidemic. (See Table ??).

We are not claiming here that closing down schools during the seasonal peak, or during an initial phase of a seasonal epidemic, will not have a beneficial effect on maximum incidence. Rather, the observed epidemiological patterns over the time period we analyzed (2003-2011) do not seem to name “school opening times” as a significant predictive factor, at least in the continental US.

### Weather data

The dataset starts with the week beginning December 31st, 2002 and includes 522 weeks (which ends exactly on the week ending December 31st, 2012). Temperature and precipitation data come from the 2.5 arcminute (approximately 4km) PRISM [46] dataset and other variables (wind speed, specific humidity, surface pressure, downward incident, and shortwave solar radiation) come from the 7.5 arcminute (approximately 12km) Phase 2 North American Land Data Assimilation System (NLDAS-2) dataset [41], [12]. These datasets are selected in large part due to the fact that both are updated in near real-time, making it possible to use these datasets for future monitoring applications. PRISM is released daily, with an approximately 13 hour delay (data for the previous day is released at 1pm EST each day) while NLDAS is released daily, with an approximately 2.5 day delay).

Variables are aggregated to county boundaries based on shapefiles from the GADM database of Global Administrative Areas [25]. Where appropriate, we considered both the average daily climate variable (for example, the daily maximum temperature averaged over the week) as well as the the maximum and/or minimum of the variable experienced over the week. For precipitation, we considered only the cumulative total precipitation experienced during the week.

### Inferring statistical ”Granger-causality” from data (Approach 1)

Granger attempted to obtain a precise definition of causal influence [28] from the following intuitive notion: Y is a cause of X, if it has unique information that alters the probabilistic estimate of the immediate future of X.

Here, we used a new, non-parametric approach to Granger causal inference [10] (Approach 1). In contrast to state-of-art binary tests [4], [32], we computed the degree of causal dependence between quantized data streams from stationary ergodic sources in a non-parametric, non-linear setting.

Our approach is significantly more general to common, regression-based implementations of Granger causal inference, and does not involve any autoregressive moving average (ARMA) modeling. All such commonly used techniques impose an a priori linear structure on the data, which then constrains the class of dependency structures we can hope to distill.

True causality, in a Humean sense, cannot be inferred [34], [35]. Among other reported approaches to ascertaining causal dependency relationships, the work of J. Pearl [47] is perhaps most visible, and builds on the paradigm of structural causal models (SCM) [48]. While Pearl’s work is often claimed to be able to answer causal queries regarding the effects of potential interventions, as well as regarding counterfactuals, our objective in this paper is somewhat different. We are interested in delineating whether infection transmission pathways can be distilled from the patterns of infection’s spatiotemporal incidence.

## Methods: Three approaches

### Analysis of quantized clinical data (Approach 1)

For the purpose of Poisson regression (Approach 2), we restricted the weeks we considered to those immediately preceding an exponential rise in incidence prevalence (see Figure 3) in order to focus on the onset of an influenza season. (We used the same consistent formulation of influenza onset across all three approaches in this study.)

**Fig. 3.**
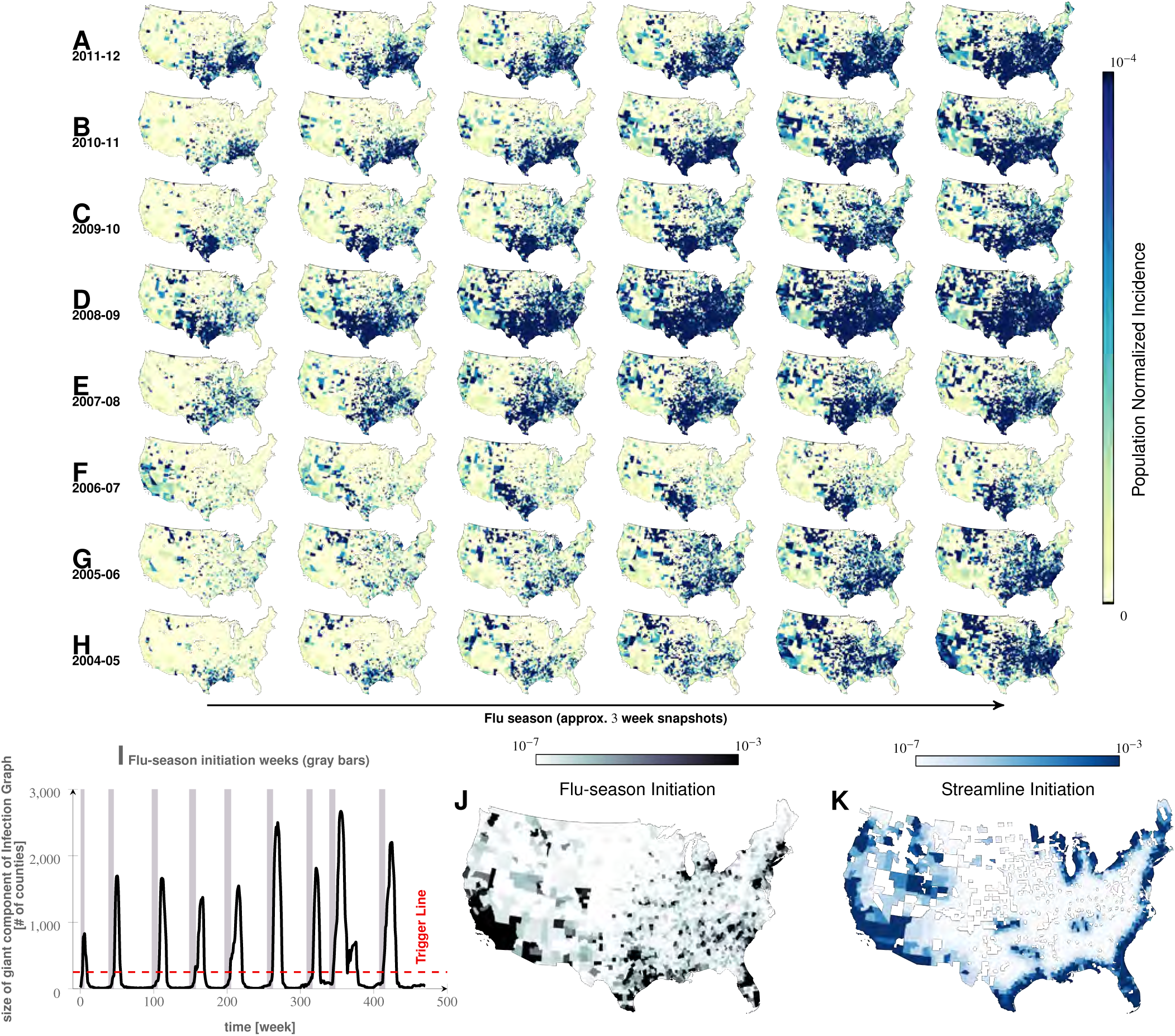
Characteristics of seasonal influenza in the continental US. An analysis of county-specific, weekly reports on the number of influenza cases for a period of 471 weeks spanning January 2003 to December 2013 (Plates A-H) for recurrent patterns of disease propagation. In particular, the weeks leading up to that in which an epidemic season peaks (determined by significant infection reports from the maximum number of counties for that season) demonstrate an apparent flow of disease from south to north, which cannot be explained by population density alone (also see movie in Supplement). Plate I illustrates the near-perfect time table for a seasonal epidemic. Plates J and K compare the county-specific initiation probabilities of an influenza season, and the causality streamlines.

For the causality analysis (Approach 1), we proceeded differently. We had an integer-valued time series for each US county, and to carry out the causality analysis, we first quantized the series in two steps:

1. Computing the difference series, i.e., the weekly change in the number of reported cases.
2. Mapping positive changes to symbol “1” and negative changes to symbol “0” (see Figure 3B).

This mapped each data series to a symbol stream over a binary alphabet. The binary quantization is not a restriction imposed by the inference algorithm; while we do require quantized magnitudes, longer data streams can be encoded with finer alphabets to accommodate an arbitrary precision. For this specific dataset, the relatively short length of the county-specific time-series necessitated a coarse quantization in order for the results to have a meaningful statistical significance.

Given a pair of such quantized streams s_a_, s_b_, the algorithm described in the Supplement computes two coefficients of causality, one for each direction. Intuitively, the coefficient
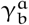
from s_a_ to s_b_ is a non-dimensional number between 0 and 1 that quantifies the amount of information that one may acquire about the second stream s_b_ from observing the first stream (s_a_). More specifically,
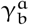
is the average reduction in the uncertainty of the next predicted symbol in stream s_b_ in bits, per bit, acquired from observed symbols in stream s_a_. It can be shown that the coefficient of causality
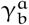
is 0 if and only if there is no causal influence from s_a_ to s_b_ in the sense of Granger, and assumes the maximum value 1 if and only if s_b_ is deterministically predictable from s_a_. Moreover,
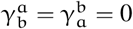
if and only if s_a_ and s_b_ are statistically independent processes [10]. It is trivial to produce examples where we would have
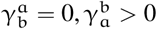
illustrating the ability of the algorithm to capture the asymmetric flow of causal influence in one preferred direction and not the other.

Additionally, whenever we computed the causality coefficient, there existed an associated notion of a time delay: We can calculate the coefficient for predicting the target stream some specified number of steps in the future, and the computed coefficient is thus parameterized by this delay. In our analysis, we computed coefficients up to a maximum delay of 10 weeks, and, for each pair of counties, selected the optimum delay which gave rise to the largest coefficient. For more details on the algorithm, see the Supplement.

### Computation of causality fields and causality streamlines (Approach 1)

To compute causality streamlines, we needed a precise notion of county neighbors. We considered two counties to be neighbors if they share either a common border, or if one county is reachable within a line-of-sight distance of 50 miles from any point within another. The latter condition removed ambiguities as to whether counties touching at a point should be still considered as neighbors. The exact distance value (50 miles) does not significantly change our results, as long as it does not vary significantly. With the definition of the neighborhood map in place, we proceeded to compute the direction-specific coefficients of causality between neighboring counties. It follows that we would obtain a set of coefficients for each county and one for each of its neighbors, capturing the degree of causal influence from a given particular county to its respective neighbors. Our algorithm also computed the probability of each coefficient arising by chance alone, and we ignore coefficients that have more than a 6% probability that two independent processes lacking any causal connection gave rise to the particular computed value of the coefficient. Once the coefficients had been calculated for each neighbor, we computed the resultant direction of causal influence outflow from that particular county. This was carried out by visualizing the causality coefficients as weights on the length of the vectors, from the centroid of the considered county to the centroids of its neighbors. We then calculated the resultant vector (see Figure 3). Viewed systematically across the continental US, these local vectors formed a discernible pattern; we can observe the emergence of a non-random structure with clearly discernible paths outlining the “causality field” (see Figure 4, Plates G, J, K). To interpret the plots, note that streamlines start at their thinnest part, their direction is indicted with thickening line; typically multiple streamlines coalesce into a river-like pattern.

**Fig. 4.**
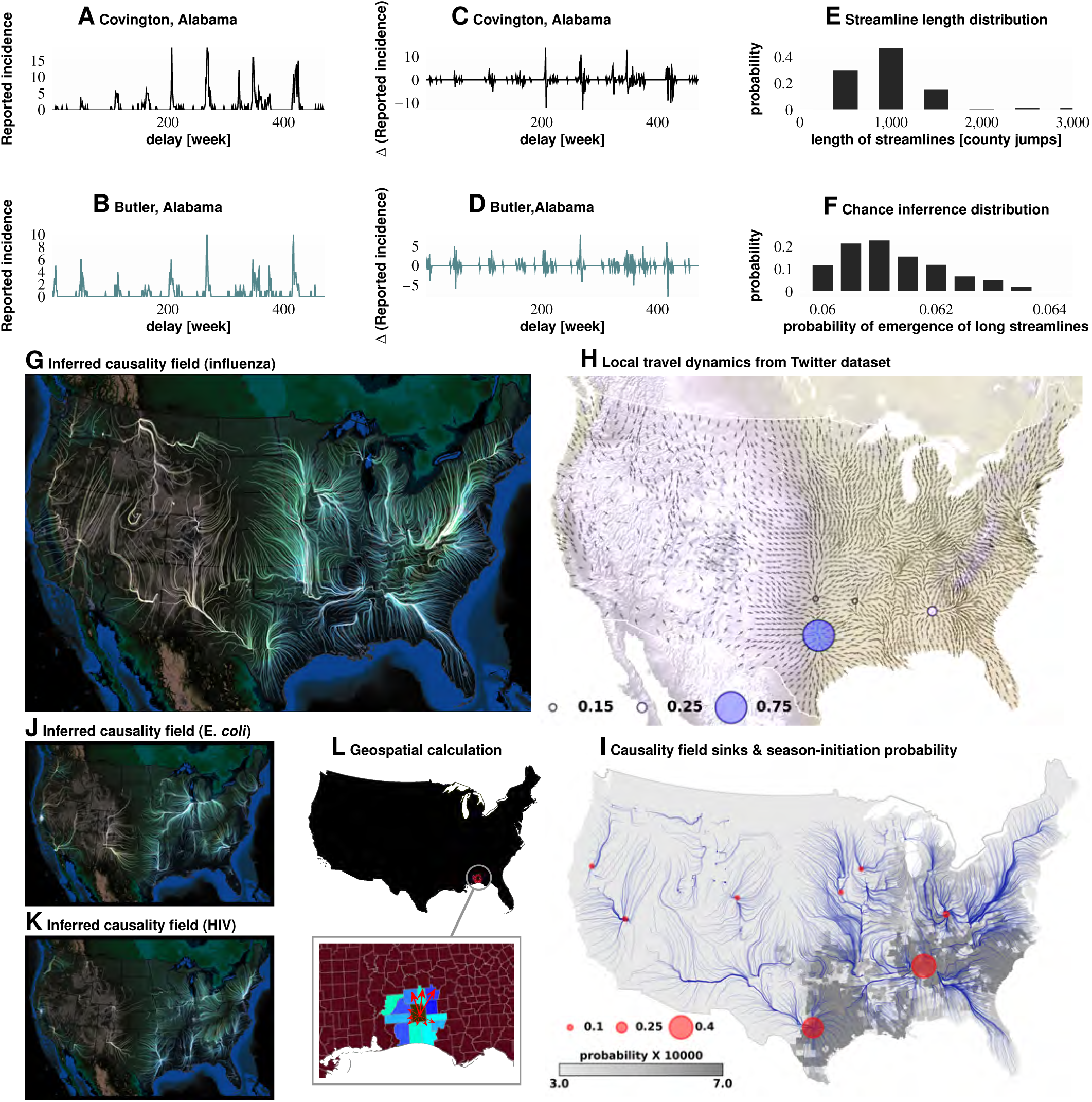
Computation of causality field, Approach 1. Plates A and B: Incidence data from neighboring counties in Alabama, US. Plates C and D: Transformation to difference-series, *i.e*., change in the number of reported cases between weeks. We imposed a binary quantization, with positive changes mapping to “1,” and negative changes mapping to “0.” From a pair of such symbol streams, we computed the direction-specific coefficients of Granger causality (see Supplement). For each county, we obtained a coefficient for each of its neighbors, which captured the degree of influence flowing outward to its respective neighbors (Plate L). We computed the expected outgoing influence by considering these coefficients as representative of the vector lengths from the centroid of the originating county to centroids of its neighbors. Viewed across the continental US, we then observed the emergence of clearly discernible paths outlining the “causality field” (Plate G). The long streamlines shown are highly significant, with the probability of chance occurrence due to accidental alignment of component stitched vectors less than 10^-185^; while each individual relationship has a chance occurrence probability of ~ 6% (Plates E and F). Plate H: Spatially-averaged travel patterns (see text in Methods) and the sink distribution between expected travel patterns. These patterns (Plate H), along with the inferred causality field (Plate I), match up closely, with sinks showing up largely in the southern US states, explaining the central role played by the Southern US. Plates J and K: Spatial analysis results for two different infections (HIV and *E. coli*, respectively) and which exhibit very different causality fields, negating the possibility of boundary effects.

### Mixed-effect Poisson regression (Approach 2)

We investigated the relative importance of putative factors as follows: Specifically, let N_ijk_ denote the total number of patients in county i, who are of age j, and gender k. Denoting the number of individuals diagnosed with influenza in a given county during given week as y_ijk_, we modeled the within-county disease incidence counts for every county (for which data was available) in the US using the following mixed-effect Poisson regression model:

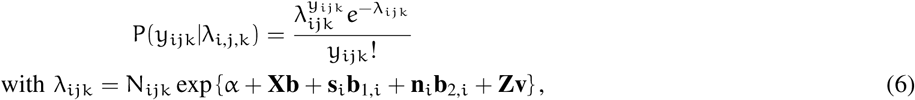

where, α is the intercept, **X** and **b** are fixed-effect design matrix and vector of fixed effects, respectively, **s**_i_ is 1 × m vector of changes in rate of infection in the i^th^ county 1,2,.., m weeks prior to the current, **b**_1,i_ is a m × 1 vector of auto-correlation fixed effects, **n**_i_ is a 1 × (3m) vector of changes in the rate of infection in the neighbors of the i^th^ county 1,2,.., m weeks prior to the current (the neighbors are subdivided into *land* neighbors, *Twitter* neighbors, and *air* neighbors), **b**_2,i_ is a (3m) × 1 vector of county-neighborhoods fixed effects, and **v** and **Z** are random effects and their design matrix, respectively. Variable m represents the depth of “memory” of auto-regression in weeks, in our case m = 4.

In this way, the total number of rows in matrix **X** is 510 × 3,143 (weeks × counties), with county-specific socioeconomic covariates and week-specific weather covariates. For disease initiation analysis, we included only a subset of time series covering approximately 50 weeks.

*Out*-*of*-*sample prediction and ROC analysis with mixed*-*effect Poisson regression:* We carried our out-of-sample prediction with the models inferred with mixed-effect regression. The steps were as follows:

1. We trained the model parameters with data from the trigger periods corresponding to the first six seasons.
2. Once the coefficients of the variables were identified, we used it to predict the response variable (influenza incidence) for the last three seasons.

As expected, the predicted incidence does not exactly match the observed out-of-sample data. Nevertheless, we see positive correlation (Figure 6, Plate A). Since we were modeling a necessarily spatio-temporal stochastic process, the predictive ability of the model is difficult to judge simply from the observed positive correlation. To resolve this, we investigated the performance of our model by computing how well it predicted the counties that experience flare-ups during the trigger-periods in the out-of-sample data. This exercise is reduced to a classification problem, by first choosing a threshold on the number of reported cases per week to define what is meant by a “flare-up” (see description below). To quantify the prediction performance, we constructed ROC curves for each of the three target seasons, for each fixed week, as follows:

1. We first quantized the incidence data to reduce it to a binary stream. In particular, we chose a threshold (10), such that for each county, and each week, we reported a “1” if the number of reported cases was greater than the threshold, and “0” otherwise. Note this quantization is different to what we used in carrying out our nonparametric Granger analysis in Approach 1.
2. For each county i, and week j, we then ended up with the binary class variable X_j,i_ ∈ {0,1} and a decision variable Y_j,i_ ∈ ℝ, where the former is the quantized incidence described above, and is the response predicted by the model, normalized between 0 and 1.
3. For a chosen decision threshold θ_D_ ∈ ℝ, we could determine the predicted class X̂_j,i_ ∈ {0,1} as:

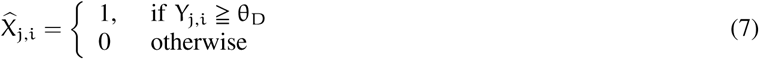
4. Comparing the observed and predicted classes, we could compute the false positive rate (FPR: defined as the ratio of false positives to the sum of false positives and true negatives), and the true positive rate (TPR: defined as the ratio of true positives to the sum of true positives and false negatives). Finally, we constructed the ROC curve, which shows the relationship of the TPR and the FPR as the decision-threshold θ_D_ is varied.
5. We constructed ROC curves for each week in the out-of-sample period, and estimated the area under curve (AUC). The AUC measures the performance of the predictor (our model) to classify correctly the counties that would go on to have a disease incidence greater than the initial set threshold (ten cases in our analysis). In the perfect case, we would have an AC of 1.0, which implies that we can achieve zero false positives, while getting a 100% true positive rate. Our best model achieves approximately 80% AUC for the trigger weeks as shown in Figure 6.

### County-matching effect analysis (Approach 3)

Let 𝕐 denote the set of all US counties, and let 𝕊 be the set of all factors we find to be significant in our mixed-effect regression analysis. Now, for any subset of factors K ⫅ 𝕊, we denote the complement set as K = 𝕊 \ K. Additionally, we define the Boolean function 𝓙: 𝕊 × 𝕐 → {true, false}, (the treatment function) where for some factor s ∈ 𝕊, and some county y e 𝕐, the Boolean value 𝓙(s, y) is **true** if the signal or treatment corresponding to the factor s is present in county y. We used he sign of the coefficients obtained in our best mixed-effect regression model to determine what counts as a positive signal. For example, because maximum specific humidity (denoted by the variable max_hus_avg) enters with a positive coefficient, if county y experiences a higher-than-average maximum specific humidity, then 𝓙(max_hus_avg,y) is true.

To simplify the notation, we used
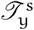
to denote a true signal, and
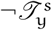
to denote a false one.

Finally, for any K ⫅ 𝕊, we define the following three sets, which we refer to as the *W*-sets:

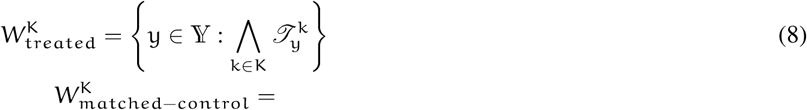

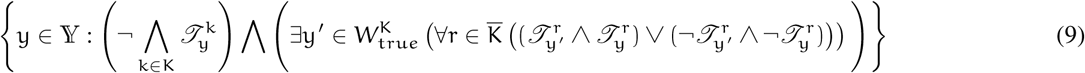

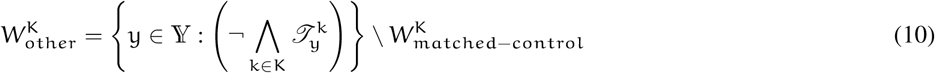

Clearly,
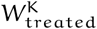
is the treated set, i.e., the set of counties which exhibit the signal encoded by the set K.
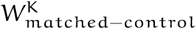
is then the matched control set of counties, which lack the signal, but each county in this set has a matching counterpart in
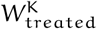.

The *W*-sets allow us to set up a 2 × 2 contingency table for any chosen subset of factors K ⫅ 𝕊 as described before. Specifically, we split the sets
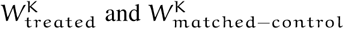
into two subsets each, representing those counties which experience a spike in influenza prevalence and which do not. The 2 × 2 contingency table is then subjected to Fisher’s exact test. The results shown in Figure 7 and Tables II and III are for one-sided tests.

**TABLE II.**
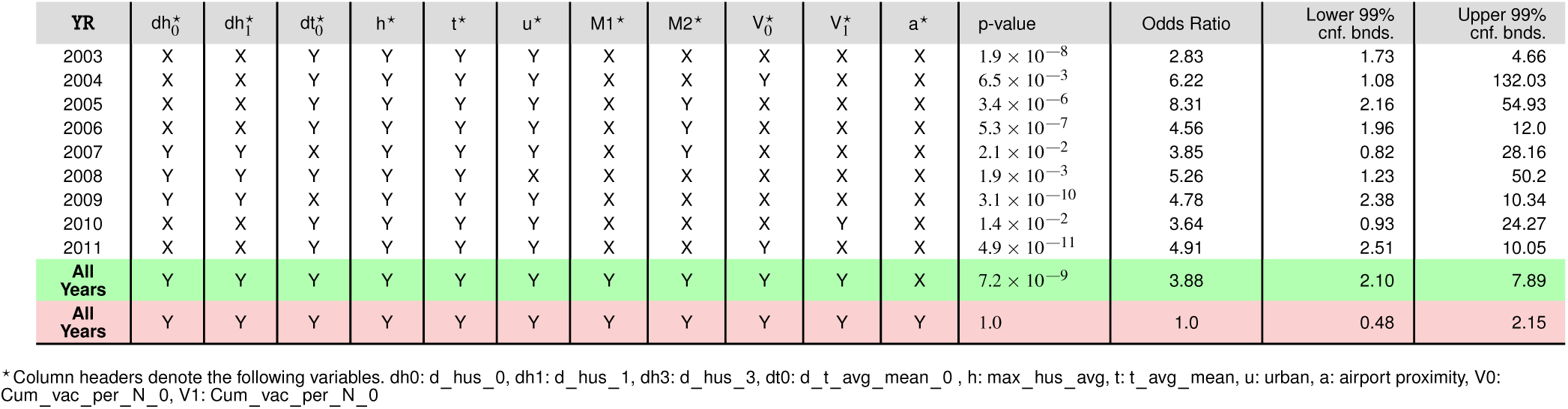
Fisher exact test results on matched treatment combinations

**TABLE III.** Fisher exact test results on matched treatment on single factors

| (a) |  |  | (b) |  |  |
| --- | --- | --- | --- | --- | --- |
| YR | p-value | 99% Conf. Bnd. | YR | p-value | 99% Conf. Bnd. |
| max_hus_avg |  |  | d_hus_1 |  |  |
| 2003 | 0.003603 | 1.0, 4.21 | 2003 | 0.1652 | 0.10, 2.43 |
| 2004 | 0.6919 | 0.16, 17.63 | 2004 | 1 | 0.22, 24.42 |
| 2005 | 0.1948 | 0.61, 7.89 | 2005 | 0.002004 | 0.09, 0.87 |
| 2006 | 0.6525 | 0.28, 3.06 | 2006 | 1 | 0.32, 7.65 |
| 2007 | 0.3574 | 0.49, 18.85 | 2007 | 0.389 | 0.33, 1.90 |
| 2008 | 0.103 | 0.55, 1.23 | 2008 | 0.02142 | 0.9, 8.48 |
| 2009 | 0.1067 | 0.68, 8.77 | 2009 | 0.1822 | 0.67, 6.06 |
| 2010 | 0.5318 | 0.27, 41.03 | 2010 | 0.6005 | 0.23, 4.22 |
| 2011 | 0.09054 | 0.74, 5.17 | 2011 | 0.9166 | 0.6, 1.88 |
| ALL YRS | $1 \times 10^{-4}$ | 1.12, 1.88 | ALL YRS | 0.07 | 0.72, 1.06 |
| t_avg_mean |  |  | d_hus_2 |  |  |
| 2003 | 0.06439 | 0.81, 3.65 | 2003 | 0.0083 | 1.01, 5.77 |
| 2004 | 1 | 0.27, 10.62 | 2004 | 0.79 | 0.36, 13.33 |
| 2005 | 0.003339 | 1.17, 123.0 | 2005 | 0.275 | 0.71, 2.54 |
| 2006 | 0.8172 | 0.29, 4.0 | 2006 | 0.24 | 0.66, 4.36 |
| 2007 | 0.537 | 0.47, 7.42 | 2007 | 0.19 | 0.62, 9.33 |
| 2008 | 0.05985 | 0.59, Inf | 2008 | 0.18 | 0.65, 7.52 |
| 2009 | 0.0006337 | 1.37, 51.68 | 2009 | 0.53 | 0.44, 6.25 |
| 2010 | 0.2853 | 0.50, 9.28 | 2010 | 0.08 | 0.21, 1.51 |
| 2011 | 0.05729 | 0.85, 3.49 | 2011 | 0.59 | 0.69, 1.87 |
| ALL YRS | $5.87 \times 10^{-9}$ | 1.36, 2.23 | ALL YRS | 0.13 | 0.78, 1.07 |
| d_hus_0 |  |  | urbanity |  |  |
| 2003 | 0.5374 | 0.55, 3.41 | 2003 | 0.0083 | 1.01, 5.77 |
| 2004 | 1 | 0.27, 11.01 | 2004 | 0.79 | 0.36, 13.33 |
| 2005 | 0.04401 | 0.81, 7.13 | 2005 | 0.275 | 0.71, 2.54 |
| 2006 | 0.001708 | 1.31, Inf | 2006 | 0.24 | 0.66, 4.36 |
| 2007 | 0.009199 | 1.0, 37.34 | 2007 | 0.19 | 0.62, 9.33 |
| 2008 | 0.3051 | 0.60, 5.92 | 2008 | 0.18 | 0.65, 7.52 |
| 2009 | 0.02726 | 0.82, 90.16 | 2009 | 0.53 | 0.44, 6.25 |
| 2010 | 1 | 0.41, 2.78 | 2010 | 0.08 | 0.21, 1.51 |
| 2011 | 0.577 | 0.57, 2.77 | 2011 | 0.59 | 0.69, 1.87 |
| ALL YRS | $1.48 \times 10^{-5}$ | 1.12, 1.64 | ALL YRS | $2.2 \times 10^{-16}$ | 3.67, 5.06 |
| d t_avg_mean_0 |  |  | airport proximity |  |  |
| 2003 | 0.004956 | 1.0, 24.12 | 2003 | 0.004956 | 1.0, 24.12 |
| 2004 | 0.445 | 0.14, 6.0 | 2004 | 0.445 | 0.14, 6.0 |
| 2005 | 0.001164 | 1.23, 12.03 | 2005 | 0.001164 | 1.23, 12.03 |
| 2006 | 0.01198 | 0.97, 11.08 | 2006 | 0.01198 | 0.97, 11.08 |
| 2007 | 0.01147 | 0.96, 11.05 | 2007 | 0.01147 | 0.96, 11.05 |
| 2008 | 0.08552 | 0.74, 11.18 | 2008 | 0.08552 | 0.74, 11.18 |
| 2009 | 0.06847 | 0.73, 17.69 | 2009 | 0.06847 | 0.73, 17.69 |
| 2010 | 0.08251 | 0.15, 1.63 | 2010 | 0.08251 | 0.15, 1.63 |
| 2011 | 0.6031 | 0.47, 3.55 | 2011 | 0.6031 | 0.47, 3.55 |
| ALL YRS | $4.98 \times 10^{-11}$ | 1.35, 2.06 | ALL YRS | $1 \times 10^{-16}$ | 1.73, 2.93 |

#### Combining factors

Infection propagation is understood to be a complex process, with multiple contributing factors (see Figures 2 and Table IV). Next, we proceed to describe findings from the three orthogonal approaches to causal analysis of the same data, then summarize their results.

**TABLE IV.** Variables in mixed-effect Poisson regression analysis (Approach 2)

(a) Definition of variables
| Variable name | physical effect |
| --- | --- |
| N | Total number of patient visits given week and county (the offset) |
| max_HUS_mean | Mean county-specific maximum specific humidity over nine years |
| d_max_HUS_i | Normalized and zero-centered deviations of maximum humidity, i=0-4 weeks before |
| t_avg_mean | Mean county-specific temperature over nine years |
| d_t_avg_i | Normalized and zero-centered deviations of mean temperature, i=0-4 weeks before |
| RSDS_mean | Mean county-specific solar insolation over nine years |
| d_RSDS_i | Normalized and zero-centered deviations of mean solar insolation, i=0-4 weeks before |
| AVG_PRESS_mean | Mean county-specific solar insolation over nine years |
| d_AVG_PRESS_i | Normalized and zero-centered deviations of mean surface pressure, i=0-4 weeks before |
| tot_prec_mean | Mean county-specific total precipitation over nine years |
| d_tot_prec_i | Normalized and zero-centered deviations of mean total precipitation, i=0-4 weeks before |
| Wind_avg_mean | Mean county-specific average wind speed over nine years |
| d_Wind_avg_i | Normalized and zero-centered deviations of average wind speed, i=0-4 weeks before |
| Income | County-specific mean income |
| airport_diffusion | Influence from proximity to airports, modeled as human traffic-weighted exponential diffusion from the 30 largest US airports |
| Am_Ind | % of American Indians in the county |
| Asian | % of Asians |
| White_Hisp | % of Caucasian/Hispanics |
| W_non_Hisp | % of Caucasian/Non-Hispanics |
| Black_Hisp | % of Black/Hispanics |
| B_non_Hisp | % of Black/Non-Hispanics |
| Pacific | % of Pacific Islanders |
| Insured | % of county population insured |
| Poor | % of county population under poverty line |
| Urban | % of county population classified as urban |
| land_i | influenza velocity change in the land neighbors of the county i weeks before the current week |
| tweet_i | influenza velocity change in the Twitter neighbors of the county i weeks before the current week |
| air_i | influenza velocity change in airport neighbors of the county i weeks before the current week |
| v_i | change in rate of infection in the county itself i weeks from the current |
| M1 | Diversity in M1 protein primary structure |
| M2 | Diversity in M2 protein primary structure |
| NA | Diversity in NA protein primary structure |
| HA | Diversity in HA protein primary structure |
| Cum_vac_per_N_i | vaccination coverage in the county cumulated over past 20 weeks i weeks from the current |

## Results

### Approach 1: Non-parametric Granger analysis

Approach 1 focuses on local and global patterns of influenza transmission: It is based on a new non-parametric, non-linear approach to Granger-causal inference. The goal of this technique is to identify infection propagation directionality, and evaluate the effects of short- and long-distance human travel on the epidemiology of influenza. Our analysis of health insurance claims covers nine years of influenza cycles (2003 to 2011, inclusively), see Figure 3. We visualize weekly, county-level prevalence as a movie (see Supplement); Figures 3 A-H show a few relevant weekly snapshots from different years. The plates in Figures 3 A-H, and especially the movie, clearly show that seasonal influenza cycles initiate in the South/Southeast US and sweep the country from south to north. This pattern repeats, with some variation, each season.

To analyze these propagation dynamics, we define the notion of a Granger *causal flow.* Treating county-specific changes in disease prevalence as a time-stamped data stream, we quantify the directional strength of the causal flow between two counties as the degree of predictability of one stream’s future, given the history of the other (see Figures 4A-D). In contrast to regression-based, classical implementations of the Granger test, our new algorithm is able to compute a *coefficient of causality:* an information-theoretic measure of the information in bits that one stream communicates about another in a direction-specific manner. A strong coefficient of this type suggests that either the disease itself, or an underlying cause, may be propagating from the former county to the latter. See the Methods section for a full discussion of Granger causal inference, including the new algorithm for non-parametric, nonlinear Granger inference.

To explore the country-wide propagation dynamics, we stitch together the properly-aligned, adjacent between-county causal flow vectors across the whole US map into *causality streamlines*, representing the aggregated spatial infection propagation over time, as shown in Figure 4G. The alignment of these flow vectors into long, continuous streamlines suggests a stable propagation mechanism across the country; the probability of a long sequence of summary movement vectors accidentally matching in the direction by mere chance is vanishingly small (p < 10^−16^ for longer streamlines).

Are there counties from which the epidemic is more likely to originate season after season? To answer this question, we follow “causality streamlines” back to their source county. Informally, influenza onset in these source counties has little or no causal dependency on their neighbors. That is, their epidemic states are seemingly caused by factors outside of disease prevalence in other counties. Figure 3K presents the county-specific likelihood of streamline initiation across our nine years of data. To verify whether this is a mere manifestation of boundary effects, we carried out identical causality analyses with different infectious disease phenotypes, specifically choosing diseases less likely to share etiologies with influenza: HIV and *Escherichia coli.* The results for both HIV and *Escherichia coli* infections are shown in Figures 4J and K, which exhibit flow patterns distinct from those obtained for influenza. These streamlines almost never originate from the coasts, thus excluding the possibility that the pattern observed for influenza is a geospatial boundary effect. Combined with the exceedingly low probability (~ 10^−185^) of chance inference for the streamlines, this strongly supports our conclusion that the epidemics are of coastal origin.

Additionally, we validated our conclusion that influenza waves tend to start in the South by directly identifying counties from which the epidemic seems to trigger. We computed a “trigger period” of five to six weeks for each season, defined as the period immediately preceding an exponential increase in influenza dispersion. To calculate this weekly dispersion, we treated each county as a node in an undirected graph, each with an edge connecting two geographically adjacent counties–only if they have both reported at least one influenza case in the specified week. We defined dispersion as the size of the largest, connected component in this undirected graph. Thus, a trigger period describes the period in which the size of the giant component of the infection graph rises above 250 counties from being under 100 as shown in Figure 3I, and then proceeds to the seasonal peak. Figure 3J presents the likelihood of a county’s being part of this largest, connected component during the trigger period. In the second approach, we followed causality streamlines back to their source county. Figure 3K presents the county-specific likelihood of streamline initiation across nine years.

These approaches produced similar results (shown in Figure 3J and Figure 3K): While epidemics seem to start in many places around the country, they successfully gain traction near large bodies of water. Otherwise, they fizzle out before triggering an actual epidemic cycle (see Figure 3J). Seasonal initiation is neither spatially uniform nor simply a reflection of county-specific population density.

Our analysis of the Twitter movement matrix indicates that people most frequently travel between neighboring counties, preferentially towards higher-population-density areas, which shows that the maximum-probability movement patterns follow the local gradient of increasing population density (see Figure S3 in the Supplement). In contrast, the geo-spatially-averaged movement vectors for each county reveal global flows in the movement patterns (see Figure 4H, along with Methods for the calculation of spatial averages).

Figures 4H-I show that average movement patterns are largely collinear to the streamline patterns.

In addition to direction of short-range travel, we used our non-parametric Granger analysis to investigate the comparative strength of short-range vs. long-range influenza propagation (see Figure 5). In the first case, we considered the neighborhood map shown in Figure 5 A (for a detailed definition of “neighbors,” see Methods), and the in the second case we considered association between major airport-bearing counties (see Figure 5 B). We then plotted the distribution of the maximum pairwise coefficient of causality, where the maximization is carried out by fixing the source and the target, and varying the delay in weeks after which we attempt to predict the target stream. It turns out that the expected value of the distribution of the coefficient is higher for short range connectivity suggesting that the short range communication of influence is stronger on average.

**Fig. 5.**
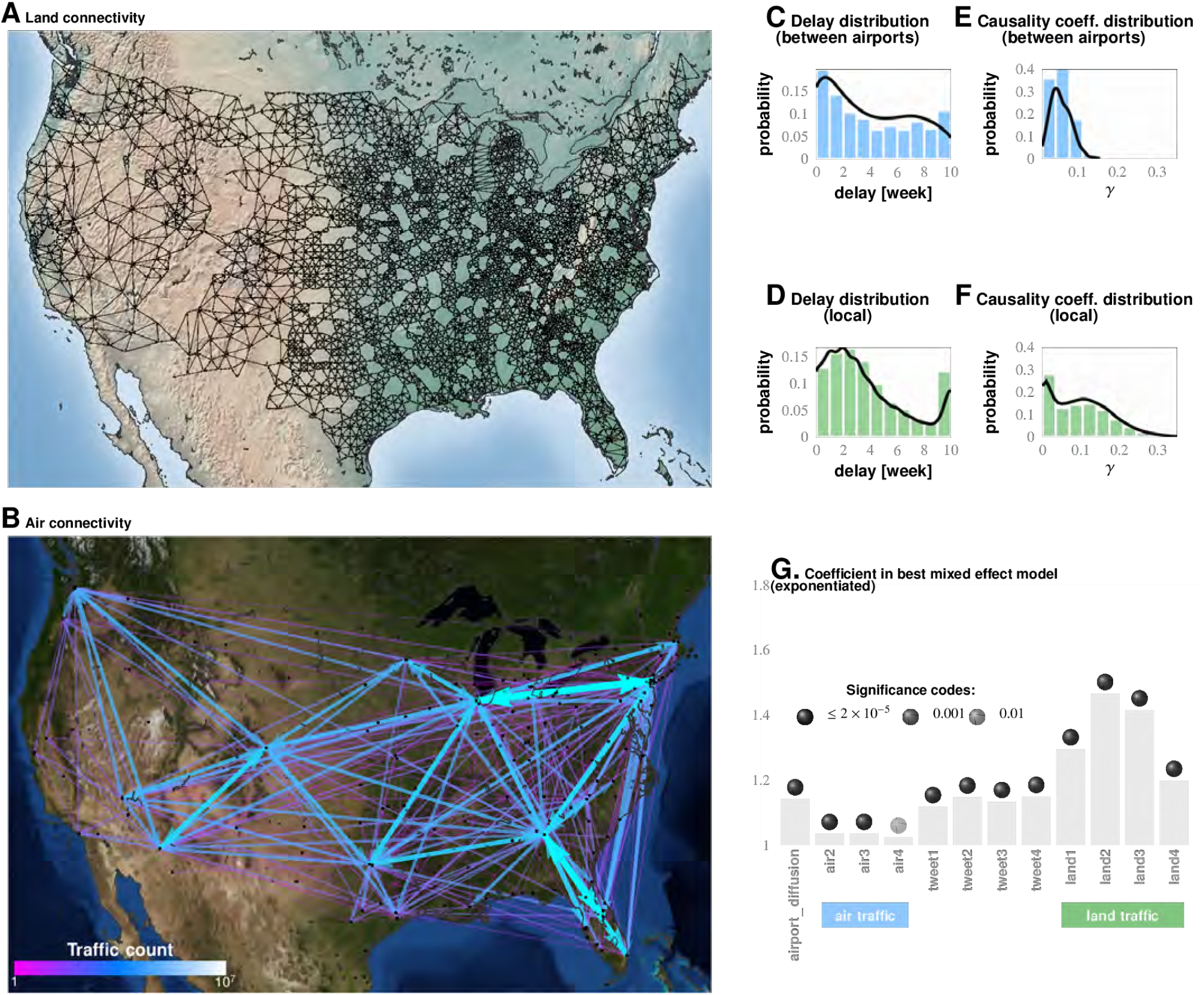
Comparing influence of short- and long-distance travel on infection propagation. Plate A shows land connectivity visualized as a graph with edges between neighboring counties. Plate B shows air connectivity as links between airports, with edge thickness proportional to traffic volume. Plate C shows the delay in weeks for the propagation of Granger-causal influence between counties in which major airports are located, and Plate E shows the distribution of the inferred causality coefficient between those same counties. Plates D and F show the delay and the causality coefficient distribution respectively, which we computed by considering spatially neighboring counties. The results show that local connectivity is more important. We reached a similar conclusion using mixed-effect Poisson regression, as shown in Plate G: The inferred coefficients for land connectivity are significantly larger than those for air connectivity, tweet-based connectivity, or exponential diffusion from the top 30 largest airports. The coefficients shown in Plate G are exponentiated, allowing us to visualize probability magnitudes (see model definition).

Conclusions associated with Approach 1: The inferred causality streamlines computed from infection time series in all counties (Figure 4) show that epidemics are mostly triggered near large water bodies, and flow inland and away. They also illustrate that the US continental southern states act as “sinks” to a large proportion of these streamlines. (“Sinks” are geographic areas that multiple streamlines converge towards; sinks are especially obvious when we look at the vector representation of causality direction. The opposite of a “sink” is a “source,” defined as an area at which at least one streamline starts.) This might explain the increased prevalence in the designated region. Additionally, the analysis shows that human travel is a very important driver of emergent epidemiological patterns, and that short-range, land-based travel is more important than air-travel. This result is cross-corroborated by our Poisson regression analysis (described next in Approach 2).

Approaches 2 and 3 are motivated by the “why” questions: (1) Why do epidemics initiate where and when they do? and; (2) Why do some disease initiations become epidemics while others do not?

### Approach 2: Importance of factors from Poisson regression

We used mixed-effect Poisson regression to investigate the identified putative factors’ relative contribution to seasonal influenza epidemic onset. Our response variable represents county-specific, weekly influenza incidence rates, segregated by gender and age. The response variable in this analysis is the total number of reported influenza patients treated in that county in a given week, using the total number of county patients treated in the same week as an offset. We used county-level random effects. All the fixed-effect variables we considered (see Table IV and the previous section) are county- and week-specific predictors, and are zero-centered and normalized by the two standard deviations. We ran a Bayesian inference of model parameters using Markov chain Monte Carlo [30].

We then focused on a subset of weeks associated with the initial rise of influenza waves (indicated by the gray bars in Figure 3I, and calculated as discussed earlier). The results from our best-fit model are illustrated in Figure 2A. We selected this particular model out of a total of 126 compared in the Bayesian analysis, a few of which we list in Table V, ranked by their decreasing goodness-of-fit, measured with the Deviance Information Criterion, DIC (See Table ?? for complete list). From the values of the inferred coefficients corresponding to the different factors, and taking into account their significance levels and credible intervals, we conclude that the roles played by weather variables, particularly humidity, appear to be significantly more complicated compared to what has been suggested in the literature.

**TABLE V:**
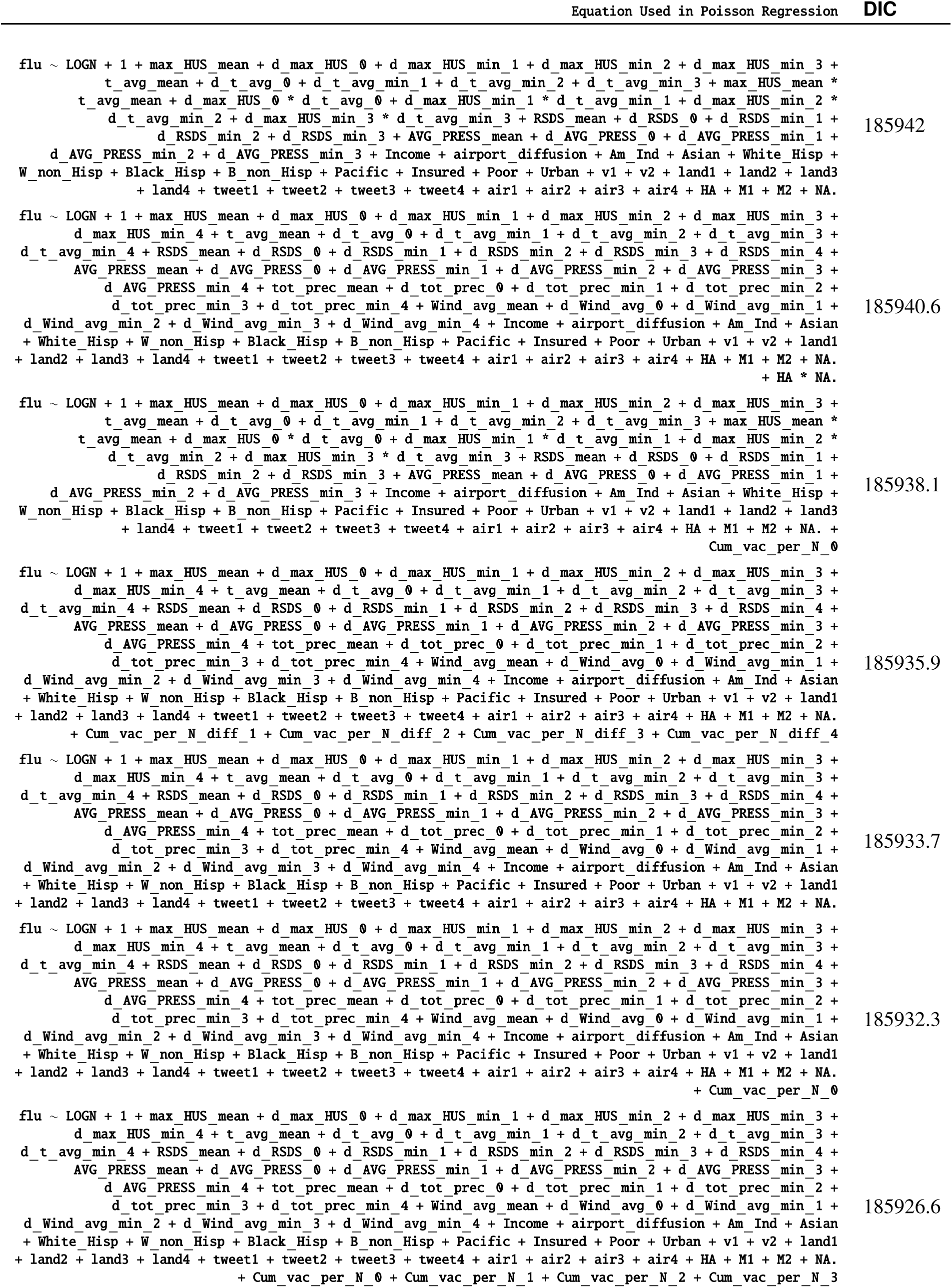
Different Models Considered, and DIC Ranking

In our analysis, we transformed all non-categorical predictor variables in our analysis by zero-centering and scaling by two standard deviations of the variable. As a result, the magnitude of regression coefficients can be directly used to judge the relative importance of individual predictors. The major conclusions of our analysis follow.

#### The surprisingly unimportant factors

School schedule was not predictive of influenza onset in our analysis (see corresponding section in Material and Methods).

Vaccination coverage failed to reach predictive significance. The variables corresponding to spatio-temporal indicators of the cumulative fraction of the inoculated population are included in our best model (see last entry in Table V), but their effect fails to be significant. However, if we drop those variables from the model, the DIC increases. We suggest that the strong dependence between antigenic diversity and vaccination coverage (see Fig. 2J-M) is responsible for this effect: Vaccination coverage is important, but its influence is captured by the antigenic variation.

#### The most important factors

The strongest predictor groups (ranked by importance) are the population’s sociodemographic properties, weather, antigenic drift of the virus, land-based travel, and autocorrelation of influenza waves.

#### Socio-demographics

The demographic make-up of the population emerges as the most important predictor, with regions with white, Hispanic, black, non-Hispanic, and Native American ethnic groups acting as probable trigger points. As showed our county-matching experiments, the level of poverty increases risk.

#### Weather

As far as weather effects are concerned, epidemics tend to originate in places with high mean maximum specific humidity, high average temperature, and low average air pressure, namely, counties at the southern, and, to a lesser extent, eastern and western US coastlines. Additionally, the spread of an epidemic is significantly influenced by a drop in specific humidity up to four weeks before its onset. However, this effect is weaker than the mean maximum specific humidity effect. Drop in average temperature that dips one to three weeks prior to the epidemic onset is also significantly important (this is consistent with earlier experiments [40]), especially when the temperature drop is accompanied by decrease in specific humidity, average wind speed, and solar flux. However, high levels of solar flux in the week of onset are also important. This complicated set of weather conditions, a signature of cold air front [53], is validated by our out-of sample predictive experiments to increase the risk of triggering the seasonal epidemic. Total precipitation also plays a positive role.

#### Is there a paradox here?

How can colder weather and lower humidity be a predictor of influenza, if influenza epidemic waves tend to start in the South with warmer climates and higher humidity? Our resolution of this seeming controversy is as follows: The stress is on the drop in humidity and temperature in those areas with high average annual values of these measurements. A temporary onset of colder, lower-humidity weather in these warm-climate areas has two effects: (1) The influenza virus can stay viable in water droplets longer than in hot, sunny weather, and; (2) Humans tend to interact indoors, in more crowded conditions. Both of these factors are favorable for transmission of the virus to the population at large.

#### Antigenic variation

Antigenic diversity for HA, NA, M1, and M2 are important predictors. While HA, NA, and M1 inhibit the trigger, M2 diversity enhances it. This peculiar difference in the direction of influence might be a manifestation of the roles played by the individual viral proteins in its life-cycle.

The first three proteins are directly involved in the viral binding to host cell surface receptors, while M2 activity is needed only during HA biosynthesis. Additionally, proteolysis experiments indicated that M2 proton channel activity helped to protect (H1N1)pdm09 HA from premature conformational changes as it traversed low-pH compartments during transport to the cell surface [1].

#### Travel

Land travel intensity one to three weeks before epidemic onset is a strong predictor. Air travel is also predictive, but its strength is an order of magnitude weaker than that of land travel.

#### Autocorrelation

The increase in influenza’s infection weekly rate one and two weeks before an epidemic onset (in the epidemic source county itself) is predictive of epidemic wave origin.

We have used substantially richer datasets than those used by earlier studies [52], [51], [55], which lends strong statistical support to our conclusions. It also allows us to disentangle and make precise the contributions from different factors, e.g., mean county humidity vs. drops in humidity before an infection. While we find the former effect to be clearly stronger (in accordance to previously reported results [52]), as are the other diverse set of factors we found to be significant.

### Validation of Predictive Capability

The robustness our results is established from different viewpoints:

1. In mixed-effect regression (Approach 2), we compare over 120 chosen model variations (see Table V for an abridged list, and Table ?? for the complete enumeration of considered models); the results appear to be qualitatively stable, i.e., while the quantitative performance of the models vary somewhat as measured by DIC for different configurations of regression equations.
2. A direct validation of predictive performance is carried out by training our model parameters for the first six seasons, and predicting the epidemic trigger locations for the last three (see Figure 6, and Materials and Methods).

**Fig. 6.**
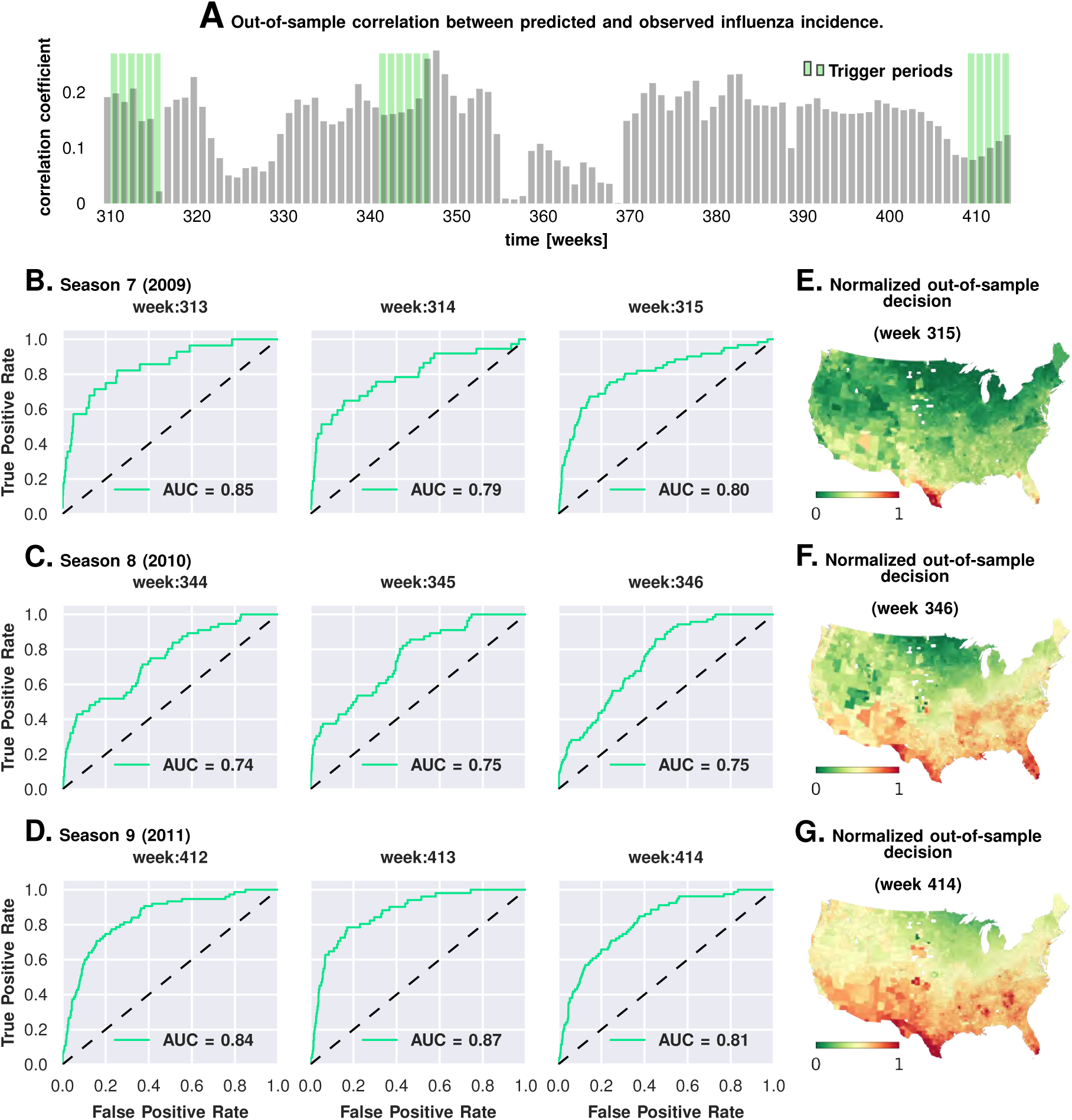
Prediction performance with training data from the first six seasons and validation on the last three. Plate A shows the correlation between the observed incidence and the model predicted response. We show significant positive correlation, particularly within the trigger periods in the out-of-sample data, which is what the model is trained for. This gives us confidence to construct ROC curves for each week. Plates B-D show the ROC curves for the last three weeks of each of the three seasons in the out-of-sample period (potentially, these computations can be repeated for all possible partitions of study weeks into training and test samples). Plates E-G illustrate that the normalized decision variable, which is the normalized response from the model, identifies the South and Southeastern counties as the trigger zones. The out-of-sample predictions of influenza incidence are always positively correlated with observed incidence (Plate A). Perhaps more importantly, we obtain good predictability as measured by the area under the curve (AUC ≈ 80%) for the receiver operating characteristics (ROC, See Plates B-D). Plates E-G show that our out-of-sample predictions correctly identify epidemic initiation in the Southern and Southeastern counties of the continental US.
3. In Approach 3 (discussed next), we conducted a corroborating matched effect analysis on the counties, using combinations of county-specific factors as a “treatment,” not unlike clinical trials in which patients on a drug regimen are matched to patients receiving a placebo [43].

### Approach 3: Matching counties & factor combinations

In the mixed-effect regression analysis (Approach 2), we jointly considered all nine epidemic seasons, leading us to conclusions that putatively apply to typical influenza dynamics in the US. It is conceivable that the significant parameter set includes multiple, partially overlapping subsets of sufficient conditions acting during individual seasons.

In Approach 3, we investigate combinations of multiple factors presented as “treatment” via a non-parametric, exact matching analysis of US counties during the weeks of epidemic onset on a season-by-season basis. For an intuitive understanding of this method, consider the effect of specific humidity on influenza prevalence. The goal of the county-matching method is to deduce associations putatively interpreted as causality relations. For example, consider testing the question of whether counties with higher-than-average mean maximum specific humidity do, indeed, have higher influenza prevalence, in a statistically significant sense, when compared to counties that do not, provided all other factors are held constant.

First, we collected the list of all counties with a drop in maximum specific humidity during the weeks leading up to an influenza season in a particular year. This is the “treated set”: the set of counties that may be thought of as subjected to the positive “treatment” of a drop in specific humidity. We split this set into two, considering counties that also experience increased influenza prevalence during the epidemic onset, and ones that do not (counties with two different values of the outcome variable). The number of counties in these two sets define the first row of a 2 × 2 contingency table. In the second row (the “control set”), we focused on counties that do not experience drop in the maximum specific humidity. However, we only considered counties that have a matching counterpart in the treated set in the following sense: For each county in the control set, we found at least one in the treated set such that the rest of the significant variables (other than specific humidity) had similar variation patterns in both counties. Once we defined the control set, we split it in the manner described for the treated set: We counted the number of control counties that experienced an increased influenza prevalence during epidemic onset, and those which did not. This defined the second row of the contingency table. Finally, we used Fisher’s exact test to compute an odds ratio (the odds of realizing these numbers by chance), along with the test-derived significance of the association between the “treatment” and epidemic wave initiation (*p*-value). Furthermore, we defined our treatment to consist of multiple factors simultaneously, e.g. specific humidity and its change in the preceding week, along with average temperature and degree of urbanity, see Figure 7.

**Fig. 7.**
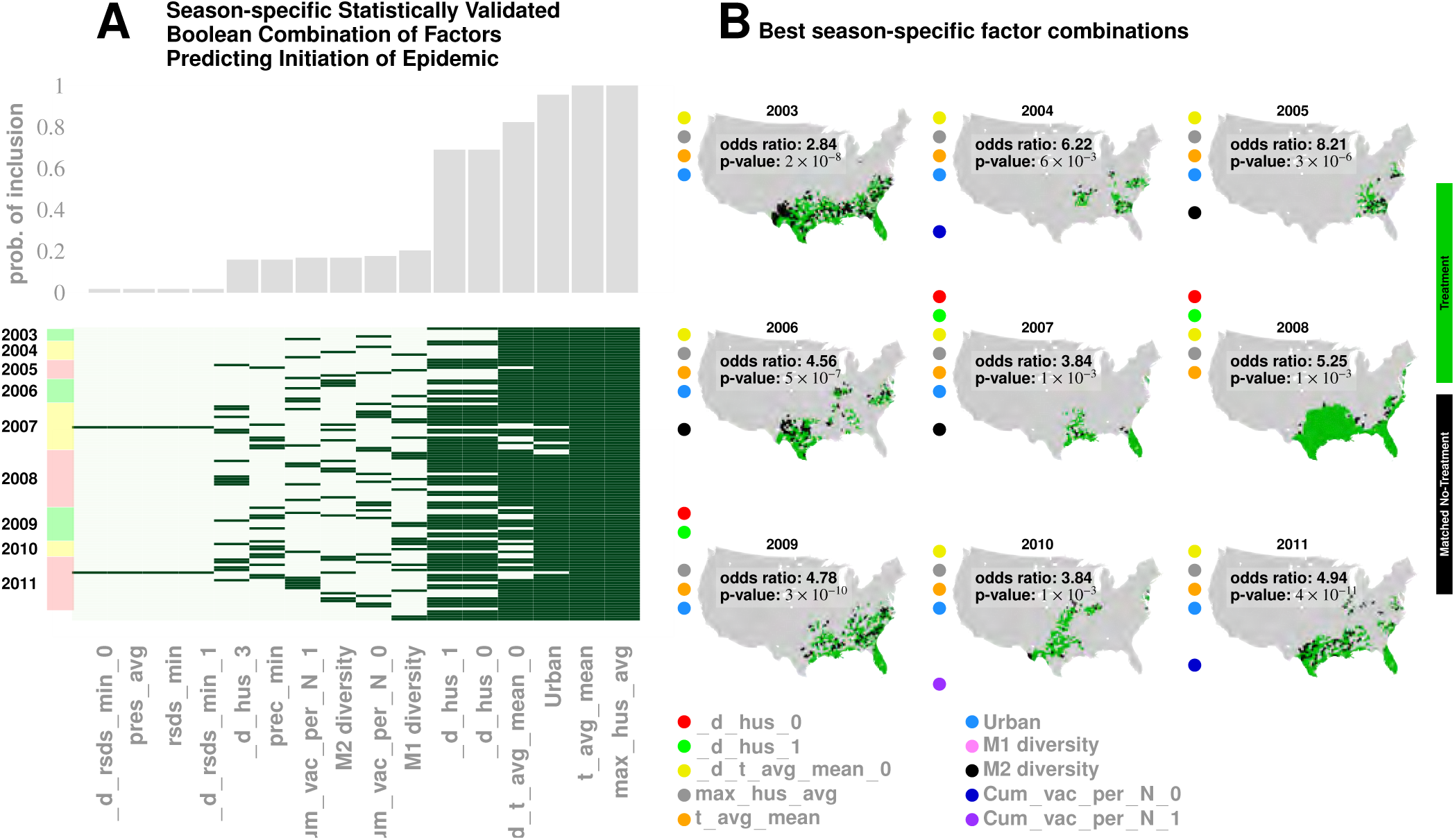
Results for our analysis involving county-matching (Approach 3). Plate A illustrates the factor combinations that turn out to be significant over the nine seasons. Notably for each season, we have multiple distinct factor sets that turn out to be significant (p < 0.05) and yield a greater than unity odd ratio. Plotting the probability with which different factors are selected when we look at season-specific county matchings (top panel in Plate A), we see a corroboration of the conclusions drawn in Approach 2. We find that specific humidity and average temperature, along with their variations are almost always included. We do see some new factors that fail to be significant in the regression analysis, e.g., degree of urbanity and vaccination coverage. While vaccination coverage is indeed included as a factor in our best performing model, in Approach 2 it failed to achieve significance, perhaps due to its strong dependence on antigenic variation (see Figure 2J-M). Degree of urbanity is indeed significant for some of the regression models we considered (see Supplementary Information), but was not significant for the model with the smallest DIC. Note that “Treatment” here is defined as a logical combination of weather factors. A treatment is typically a conjunction of several weather variables. For example, the treatment shown in top left panel of Plate B involves a conjunction of: (1) a drop in average temperature during the week of infection; (2) a drop in temperature during the week of infection; (3) a higher-than-average specific humidity; (4) a higher-than-average temperature, and; (5) a high degree of urbanity. With respect to the “treatment,” we can divide counties into three groups: (1) “treated counties,” shown in green; (2) at least one matching county for each of the treated counties (matching counties are very close to the treated counties in all aspects but in treatment, which we called “control” counties), shown in black, and; (3) other counties, shown in grey. The counties in the “treatment” and “control” groups are further subdivided into those counties that initiated an influenza wave and those that have not, resulting in four counts arranged into a two-by-two contingency table. We then used the Fisher exact test to test for association between treatment and influenza onset. Panels in Plate B show both the treated and control sets for the 9 seasons for a subset of chosen factors. The results are significant, as shown in Tables II and III. The variable definitions are given in Table IV. Notably, some of the variables found significant in the regression analysis are not included above, and some which are not found to be significant in the best regression model show up here. This is not to imply that they are not predictive or lack causal influence. The matched treatment approach, as described above, is not very effective if we use more than ~ 10 – 15 factors simultaneously to define the treated set (for the amount of data we have); this results in a contingency table populated with zero entries.

Unlike the mixed-effect regression approach, this matched analysis is non-parametric, and should reveal whether multiple factors are, indeed, simultaneously necessary. More importantly, it should reveal whether the previous results are somehow artifacts of our chosen, specific model structures.

The results of Approach 3’s analysis are presented in Figure 7 and Tables II and III. We find that no single variable is able to consistently yield a statistically significant odds ratio greater than one; multiple factors interact to shape an epidemic trigger (see Table III for a few examples). With a total of 47 significant variables in our best mixed-effect model, an exhaustive search for all combinations is not feasible. Instead, we performed a standard evolutionary search, looking for combinations that yielded a significant odds ratio for individual seasons. Additionally, we considered all seasons together (by simply adding the contingency tables, element-wise) in order to increase the test’s statistical power.

We isolated ten variables (as shown in Figure 7, Plate B) in this manner which included maximum specific humidity and average temperature along with their variations, the degree of urbanity, antigenic variation, and vaccination coverage.

The factors that appeared most often in our analysis are illustrated in Plate A: It appears that maximum specific humidity and average temperature, along with their variations, and the degree of urbanity have the most frequent contribution, followed by antigenic variation and vaccination coverage.

We do see some new factors here that fail to be of significance in the regression analysis (Approach 2), e.g., degree of urbanity and vaccination coverage. While vaccination coverage is included as a factor in our best performing model in Approach 2, it failed to achieve significance, perhaps due to its strong dependence on antigenic variation (see Figures 2J-M). Degree of urbanity is indeed significant for some of the regression models we considered (see Supplementary Information), but failed to be so for the model with the least DIC.

Thus, Approach 3, while not a reflection of the conclusions from Approach 2, corroborates and strengthens its key claims.

The exact set of factors vary somewhat over the seasons; nevertheless, they together yield significant results when all seasons are considered together. Airport proximity, when considered as the sole driving factor (the sole treatment), fails to yield significant odds ratio. It also fails to to achieve significance when acting in conjunction with the remaining nine factors. The second, crucial conclusion corroborates our conclusion from both the mixed-effect regression and the geographic streamline analyses: The sets of counties treated are near coasts on the southern region of the continental US (see Plates A - I in Figure 7).

## Discussion

The following aspects make our study of influenza triggers new in the influenza literature: (1) Instead of simulating the plausibility of one particular epidemic trigger model with a dynamic disease transmission model, we used formal model selection tools to compare goodness of fit of hundreds of plausible models; (2) We explicitly attempted to systematically cross-compare the importance of numerous individual factors typically hypothesized to contribute to epidemic onset; (3) To accomplish this, we collected an unprecedented volume of temporal and spatial data on disease dynamics and the dynamics of putative predisposing factors; (4) We used several orthogonal computational causality-inference techniques (one of which was developed specifically for this study) to probe associations between disease onset and putative epidemic triggers; (5) We tested our best models for their predictive potential and demonstrated that they are, indeed, suitable for forecasting disease waves, and; (6) For the first time, numerous candidate factors combined in a single, integrative study.

Convergent conclusions, culled from such radically different techniques, strengthen our claims, and make it statistically unlikely that we are observing analysis artifacts. First, the Granger causality analysis results (Approach 1) provide insights into the details of influenza’s epidemiological dynamics. Figure 4G traces out the paths most likely followed by the infection, on average, across the continental US. We note that ~ 75% of the streamlines sink in counties belonging to the southern states, which matches up well with the average prevalence over nine years (see Figure 4I). What drives this particular causality field’s geometry? While we cannot definitively answer this question, a comparison of the global patterns emerges from the local mobility data culled from the aforementioned Twitter database and offers a tentative explanation (see Figure 4H). Secondly, contrary to human travel patterns’ reported influence on seasonal epidemics, [59] (but consistent with [26]), we find that short-distance travel contributes more significantly to disease spread (see Figure 5). In particular, we find that long-range air travel is important as an epidemic *trigger*, but once infection waves are triggered, air travel patterns (or proximity to major airports) become less important. Short-range mobility, on the other hand, is apparently important for sustaining infection transmission over each season. Thus, we find short-range travel to be more important for defining the emergent spatio-temporal geometry of infection waves, while proximity to airports is more important for actually triggering an influenza season; the latter loses positive influence once an infection is under way. This conclusion is justified as follows: 1) Employing regression calculations using all weeks, as opposed to limiting ourselves to the few weeks before a season’s onset, results for airport proximity having a statistically significant negative influence (see Supplement). 2) Results from our Granger-causal inference indicate that, on average, the local putatively causal connections are far stronger compared to the putatively causal connections between counties within which the major airports are located. (see Figure 5 C and F). Additionally, from our best mixed-effect regression model (Figure 2A), we find that land connectivity effects are significantly stronger than air connectivity effects. The predictive value of Twitter connectivity, which intuitively captures both local and long-distance travel, lies in-between land and air connectivity coefficients. Note that Twitter connectivity is represented as a directed graph, where for each pair of counties, i and j, the (i, j) edge weight represents the conditional probability of ending at county j, given that a traveler/Twitter user started her journey in county i. Transition probabilities from i to j sum to 1 over all j. Therefore, intuitively, the Twitter connectivity graph should have the features of both a land-connectivity and an air travel graph; which indeed appears to be close to reality. 3) Finally, while airport diffusion is a significant factor in our best Poisson model (using data from the initiation period), the causal streamlines (constructed with the complete, all-year incidence data) do not seem to originate from airport-bearing counties.

The role of short-distance travel is particularly crucial in explaining influenza’s time-averaged geo-spatial prevalence. While the mixed-effect regression analysis explains seasonal initiation in the vicinity of the continental US southern shores, it might not, by itself, adequately explain its average prevalence patterns across the country.

Also not explained solely by our regression models is the occurrence of relatively high infection prevalence in the central parts of the country. These differences cannot be attributed to long-distance air travel, as discussed before. However, the routes taken by the causality streamlines (as computed by the non-parametric Granger analysis), interpreted as paths followed by an infection on average, suggest an explanation: The close match between the Granger-causal flow and the short-range mobility patterns (derived from Twitter analysis) strongly suggest that average disease prevalence is modulated by short-range mobility.

A summary of the complex relationship between the driving factors that contribute to the trigger, and, subsequently, to the development of a seasonal epidemic can be clarified with a forest fire methaphor.

The maturation of a forest fire requires the collusion of multiple factors–namely the presence of flammable media, the initiating spark, and a wind current to help to spread the fire. Our conceived mapping of this analogy to influenza infection is as follows:

1. Flammable Media: The Southern US appears to have an unusually high level of social connectivity; it is at least one order of magnitude higher than that in the north of the country (see GSS survey results [54] and Table I). The number of close friends, close friends who are neighbors, and communities of people who all, or mostly, know each other is much higher in the South than in the country at large. Our conjecture is that a manifestation of this high-connectivity is the highest percentage of people infected with influenza (20%) at the peak of infection (as opposed to 4% in other parts of the country).
2. Initiation Spark: An initial spark for the infection wave is generated by a combination of weather and demographic factors. Specifically, warm, humid places are conducive to influenza wave initiation - particularly in weeks where specific humidity drops. Nearness to airports is also important, as well as demographic and economic makeup, and also the degree of urbanization. Note that this first static condition (warm humid places) is highly correlated with areas in the South with greater social connectivity. It is possible that static meteorological variables (warm mean temperature and high mean humidity) serve as proxies for high social connectivity or other correlated socioeconomic factors.
3. Wind: The “wind” in this analogy is the collective movement of a large number of people, integrated over time, revealing persistent “currents.” These currents reproducibly point from coastlines and move inwards towards the center of the continent, making them perfect vehicles to transmit the infection inland from the shores.

Each of our three types of computational approaches has their strengths and weaknesses: (1) The Poisson mixed-effect regression allows for the direct comparison of the predictive strength of numerous predictor variables and accounts for spatial and temporal autocorrelation, but relies on strong modeling assumptions; (2) Though the non-parametric Granger analysis is not limited by restrictive modeling assumptions, in our implementation, it focuses only on trends of infection propagation between counties, and; (3) The county-matching analysis is also model-free, but this freedom comes at the expense of lesser statistical power.

We conclude by highlighting the structure of overlapping conclusions delivered by our three approaches, (see Figure 1 Plate A). Approach 1: Granger-causality analysis suggests that an epidemic tends to begin in the South, near water bodies, that short-range, land-based travel is more influential compared to air-travel for infection propagation, and provides a map of mean infection flow across the continental US. Approach 2: Poisson regression identifies significant predictive factors, ranks these factors by importance, suggests that Southern shores are where the epidemic begins, and corroborates Approach 1’s result on short-range vs. long-range travel. Approach 3 (county-matching) further narrows down epidemic onset source to the Southeastern shores of continental US, and identifies a smaller validated subset of predictive factors.

## Acknowledgements

We thank Erin Gannon and Margarita Rzhetsky for numerous comments on earlier versions of the manuscript. This work was supported by NIH grants 1P50MH094267, U01HL108634-01, R01HL122712-02, R01GM100467, and U01GM110748, in addition to DARPA contract W911NF1410333, and a gift from Liz and Kent Dauten.

